# Homozygous ALS-linked mutations in TARDBP/TDP-43 lead to hypoactivity and synaptic abnormalities in human iPSC-derived motor neurons

**DOI:** 10.1101/2023.03.22.533562

**Authors:** Sarah Lépine, Angela Nauleau-Javaudin, Eric Deneault, Carol X.-Q. Chen, Narges Abdian, Anna Krystina Franco-Flores, Ghazal Haghi, María José Castellanos-Montiel, Gilles Maussion, Mathilde Chaineau, Thomas M. Durcan

## Abstract

Cytoplasmic mislocalization and aggregation of the RNA-binding protein TDP-43 is a pathological hallmark of the motor neuron (MN) disease amyotrophic lateral sclerosis (ALS). Furthermore, while mutations in the *TARDBP* gene (encoding TDP-43) have been associated with ALS, the pathogenic consequences of these mutations remain poorly understood. Using CRISPR/Cas9, we engineered two homozygous knock-in iPSC lines carrying mutations in *TARDBP* encoding TDP-43^A382T^ and TDP-43^G348C^, two common yet understudied ALS TDP-43 variants. MNs differentiated from knock-in iPSCs had normal viability and displayed no significant changes in TDP-43 subcellular localization, phosphorylation, solubility, or aggregation compared with isogenic control MNs. However, our results highlight synaptic impairments in both TDP-43^A382T^ and TDP-43^G348C^ MN cultures, as reflected in synapse abnormalities and alterations in spontaneous neuronal activity. Collectively, our findings suggest that MN dysfunction may precede the occurrence of TDP-43 pathology and neurodegeneration in ALS and further implicate synaptic and excitability defects in the pathobiology of this disease.

## Introduction

Amyotrophic lateral sclerosis (ALS) is a neurodegenerative disorder characterized by the progressive loss of motor neurons (MNs) in the brain and the spinal cord, resulting in weakness and paralysis that is usually fatal within two to four years after onset.^1^ While about 10% of ALS cases follow a pattern of inheritance (termed familial ALS (fALS)), the majority of cases occur in the absence of a clear family history (sporadic ALS (sALS)). Overall, it is estimated that 15-20% of cases have a known genetic cause.^2^ The *TARDBP* gene (encoding TDP-43) is among the most commonly mutated ALS-associated genes after *C9ORF72*, *SOD1* and *FUS*, with nearly 40 missense mutations identified in patients to date accounting for ∼3% (fALS) and <1% (sALS) of cases^3^ (ALSoD database; https://alsod.ac.uk/). At the neuropathological level, the cytoplasmic mislocalization and aggregation of TDP-43 is a signature feature of ALS. These pathological changes, known as TDP-43 pathology, are observed in post-mortem tissues of >95% of patients,^4–6^ suggesting that convergent mechanisms of TDP-43 dysfunction are involved in both familial and sporadic disease. Thus, identifying the mechanisms through which TDP-43 dysregulation contributes to disease pathogenesis is of foremost importance in developing new therapeutics for ALS.

TDP-43 is a DNA/RNA binding protein involved in several steps of RNA processing including transcription, splicing, RNA transport, and translation.^7–10^ Early efforts to decipher the pathological roles of TDP-43 in ALS have primarily focused on overexpression and loss-of-function models and demonstrated that TDP-43 levels must be tightly regulated for it to exert its normal cellular functions. Indeed, genetic ablation of *TARDBP* is lethal during embryogenesis,^11,12^ and acute TDP-43 depletion (i.e., via conditional knockout or RNA interference) leads to neurodegeneration and ALS-like manifestations in mice.^13–15^ Similarly, overexpression of ALS-associated TDP-43 variants or even the wild-type protein exerts deleterious effects across species, including motor deficits, shortened lifespan, and MN loss in animal models,^16–20^ and cytotoxicity and various cellular dysfunctions in human immortalized neuron-like cells.^21–23^ Given the potential confounding effects of overexpression strategies, discerning the pathological contributions of *TARDBP* mutations has been challenging.

Progress in induced pluripotent stem cell (iPSC) technology now offer an unprecedented opportunity to study the impact of disease-associated mutations in human disease-relevant cell types. However, a common difficulty in studying rare mutations is the recruitment of patients and access to their cells for research. Taking advantage of the CRISPR/Cas9 technology, mutant iPSCs can be generated by introducing a mutation of interest into the genome of a control (wild-type) iPSC line. Our group and others have established robust workflows for quality control, gene editing, and differentiation of iPSCs into several cell types, including MNs.^24–27^

In recent years, a number of iPSC-derived models have been generated to assess the effects of *TARDBP* mutations expressed at endogenous levels *in vitro* (reviewed by Hawrot et al^28^). Although both gain- and loss-of-function mechanisms have been proposed, the pathogenic properties of ALS TDP-43 variants remain poorly understood. In particular, the pathologic manifestations of the TDP-43^A382T^ and TDP-43^G348C^ variants - the first and third most frequent ALS variants of TDP-43, respectively (ALSoD database; https://alsod.ac.uk/) - have not yet been fully characterized in a human model system.

To address this gap, we utilized CRISPR/Cas9 to generate two homozygous knock-in iPSC lines carrying point mutations in *TARDBP* coding for TDP-43^A382T^ or TDP-43^G348C^. We found that these mutations did not cause overt neurodegeneration nor TDP-43 aggregation or cytoplasmic mislocalization. Furthermore, mutant MNs did not recapitulate other biochemical hallmarks of pathologically altered TDP-43, including phosphorylation, C-terminal cleavage, and accumulation of detergent-insoluble species. Despite the apparent absence of TDP-43 pathology, our results highlight synaptic abnormalities and decreased neuronal activity in mutant MNs, pointing to synaptic dysfunction as an early event in ALS pathogenesis.

## Results

### Generation of *TARDBP* knock-in iPSCs lines using CRIPSR/Cas9

The vast majority of *TARDBP* mutations cluster in exon 6 of the gene, which encodes the protein’s C-terminal domain. Using CRISPR/Cas9 technology, we edited a well characterized healthy control iPSC line (i.e., AIW002-02)^26^ to generate two homozygous knock-in *TARDBP* iPSC lines expressing TDP-43^A382T^ or TDP-43^G348C^ (**Table S1**). Gene editing was performed by nucleofection of (i) the Cas9 nuclease, (ii) the single guide RNA (sgRNA) targeting the edit site of *TARDBP* (exon 6), and (iii) the single-stranded donor oligonucleotide (ssODN) carrying the mutation and the homology arms to enable integration into the genome via homology-directed repair (**Figure 1A**; **Table S2**). Successful introduction of the mutations and homozygosity of the iPSC lines were confirmed by digital droplet PCR (ddPCR) and Sanger sequencing (**Figures 1B** and **S1A-S1B**; **Table S3**). The pluripotency of gene-edited iPSCs was verified by performing immunocytochemistry for pluripotency-associated markers Nanog, TRA-1-60, SSEA-4, and Oct-3/4 (**Figure S2B**). Genome stability testing confirmed that *TARDBP* knock-in iPSC lines maintained normal karyotypes and chromosome copy numbers (**Figure S2C-S2D**). Isogeneity with the parental control iPSC line was verified using short-tandem repeat (STR) profiling (**Figure S2E**).

**Figure 1.**
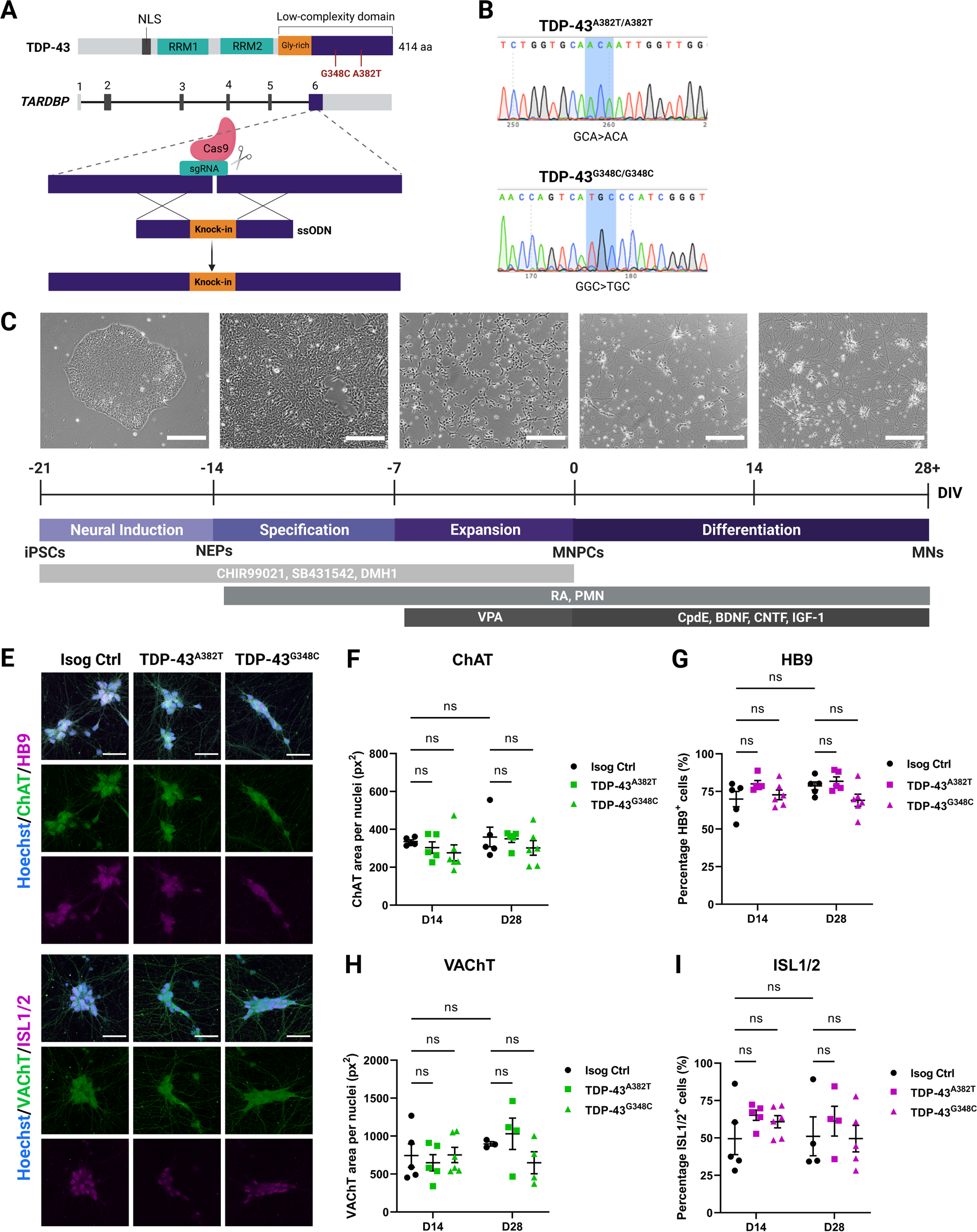
Generation of *TARDBP* knock-in iPSC lines and differentiation into MNs. (A) Schematic representation of CRISPR/Cas9-mediated genome editing via homology-directed repair. (B) iPSC lines genotyping using Sanger sequencing. (C) Schematic representation of the protocol for sequential differentiation of iPSCs into neuroepithelial progenitors (NEPs), MNPCs, and MNs with representative phase-contrast images of cells along differentiation. Scale bar, 250 μm. For time-lapse movie depicting maturation of MNPCs into MNs, see Video S1. (E-I) Representative images (E) and quantification (F-I) of MNs differentiated for 2 weeks (D14) and 4 weeks (D28) subjected to immunocytochemistry for the common MN markers HB9, ISL1/2, ChAT and VAChT. Scale bar, 50 μm. Data shown as mean ± SEM. n=5 independent experiments. See also Figures S1 to S4.

### *TARDBP* mutations do not impair normal differentiation of iPSCs into MNs

The two knock-in iPSC lines (TDP-43^A382T^ and TDP-43^G348C^) and the AIW002-02 parental isogenic control line were differentiated into MNs using a previously published protocol that mimics MN differentiation during development (**Figure 1C**).^25^ Cells were harvested at multiple timepoints to characterize the differentiation process and validate their identity. iPSCs were initially induced into neuroepithelial progenitors (NEPs) via dual-SMAD signaling inhibition, followed by specification into MN progenitor cells (MNPCs). MNPCs showed immunoreactivity for progenitor markers OLIG2, PAX6, and Nestin without significant differences between mutant and isogenic control cultures (OLIG2 TDP-43^A382T^: p=0.4780; TDP-43^G348C^ p=9062; PAX6 TDP-43^A382T^: p=0.1372; TDP-43^G348C^: p=0.0689; Nestin TDP-43^A382T^: p>0.9999; TDP-43^G348C^: p=0.1794 by one-way ANOVA) (**Figure S3A-S3D**). MNPCs were then cryopreserved or plated for final differentiation into MNs (**Video S1**). After two- and four-weeks post-plating of MNPCs, immunocytochemical analyses revealed no difference in the proportion of HB9^+^ and ISL1/2^+^ MNs (Two-weeks post-plating: HB9 p=0.1091 (TDP-43^A382T^), p=0.7932 (TDP-43^G348C^); ISL1/2 p=0.3162 (TDP-43^A382T^), p=0.4999 (TDP-43^G348C^). Four-weeks post-plating: HB9 p=0.7683 (TDP-43^A382T^), p=0.1141 (TDP-43^G348C^); ISL1/2 p=0.6579 (TDP-43^A382T^), p=0.9883 (TDP-43^G348C^)) or expression of cholinergic markers ChAT and VAChT (Two-weeks post-plating: ChAT p=0.7677 (TDP-43^A382T^), p=0.3992 (TDP-43^G348C^); VAChT p=0.8262 (TDP-43^A382T^), p=0.9985 (TDP-43^G348C^). Four-weeks post-plating: ChAT p=0.9769 (TDP-43^A382T^), p=0.4185 (TDP-43^G348C^); VAChT p=0.7693 (TDP-43^A382T^), p=0.4318 (TDP-43^G348C^)) among mutant and isogenic control cultures by two-way ANOVA. (**Figure 1E-1I**). Further tests showed statistically equivalent expression of MN markers to isogenic control MNs by four-weeks post-plating, except for HB9 and VAChT in TDP-43^G348C^ MNs compared with isogenic control (See **Figure S4A-S4C** and **STAR Methods** for equivalence testing). Quantitative PCR (qPCR) analysis examining longitudinal transcript levels of developmental markers confirmed that mutant and isogenic control cultures downregulated MNPC markers and upregulated MN markers as they differentiated into MNs (**Figure S3E**). Additionally, differentiating MNs upregulated *FOXP1* and downregulated *LHX3* (**Figure S3E**), consistent with limb-innervating lateral motor column (LMC) MN identity.^29^ LMC MNs (*FOXP1*^+^/*LHX3^−^*) are most susceptible to neurodegeneration in the majority of ALS patients, where disease typically first manifests by focal weakness in distal limb muscles.^30^ Lastly, immunostaining with two neuronal markers (NF-H and βIII-tubulin) revealed that TDP-43 MN cultures formed a dense axonal network with similar morphological features relative to control conditions, with no differences in total axonal area and branching (**Figure 2A-2D**). Levels of NF-H and βIII-tubulin, further examined by western blotting, did not significantly differ between conditions (NF-H TDP-43^A382T^: p=0.8401; TDP-43^G348C^: p=0.5045; βIII-tubulin TDP-43^A382T^: p=0.9269; TDP-43^G348C^: p=0.7456 by one-way ANOVA) (**Figure S3F-S3H**). Overall, these results indicate that TDP-43^A382T^ and TDP-43^G348C^ do not impair the normal differentiation of iPSCs into MNs.

**Figure 2.**
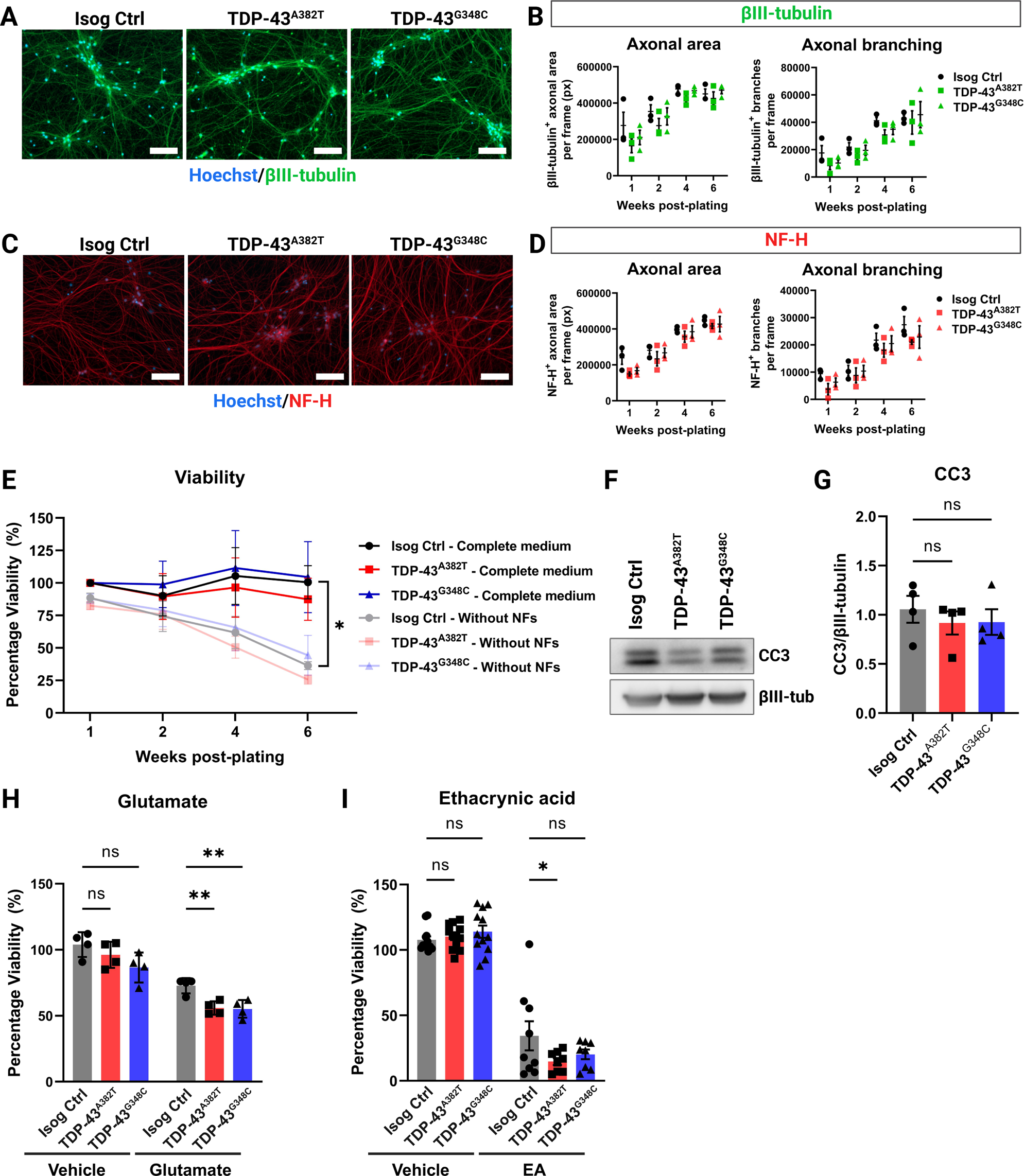
TDP-43 MN cultures form a normal axonal network and maintain viability. (A,C) Representative images of MNs differentiated for 6 weeks subjected to immunocytochemistry for neuronal markers βIII-tubulin (A) and NF-H (C). Scale bar, 100 µm. (B,D) Quantification of total area and number of branches of βIII-tubulin^+^ axons (B) and NF-H^+^ axons (D). n=3 independent experiments. (E) Viability of MN cultures differentiated with and without neurotrophic factors (NF) supplementation over a span of 6 weeks post-plating. n=4 independent experiments. (F-G) Immunoblot (F) and quantification (G) of cleaved caspase 3 (CC3) levels. βIII-tubulin was used as loading control. Extractions were performed in MNs harvested after 6 weeks post-plating. n=4 independent experiments. (H) Effect of glutamate treatment (0.1 mM glutamate, 24 h) on viability of MNs differentiated for 4 weeks. n=4 independent experiments. (I) Effect of ethacrynic acid treatment (50 μM EA, 17 h) on viability of MNs differentiated for 4 weeks. Individual points represent per-well values from 3 independent experiments. All data shown as mean ± SEM. \**p*<0.05, \*\**p*<0.01, \*\*\**p*<0.001. See also Figures S3 and S4.

### TDP-43 MN cultures maintain viability but are more susceptible to cellular stress

Neurodegeneration is a core feature of ALS. For this reason, we examined whether MNs derived from *TARDBP* knock-in iPSCs were more vulnerable to cell death. We analyzed the viability of MNs differentiated for 2, 4 and 6 weeks using an ATP-based luminescent viability assay (**Figure 2E**). TDP-43 MN cultures survived at comparable levels to the isogenic control under basal conditions (6 weeks timepoint TDP-43^A382T^: p=0.9880; TDP-43^G348C^: p>0.9999 by two-way ANOVA; see **Figure S4D-S4F** for equivalence testing) (**Figure 2E**). As caspase activation can occur before detectable neuron loss,^31,32^ we also examined levels of the apoptosis marker cleaved caspase 3 (CC3) at 6-weeks post-plating using western blotting (**Figure 2F**). In concordance with viability assays, CC3 levels did not significantly differ between the mutant and control cultures (TDP-43^A382T^: p=0.6739; TDP-43^G348C^: p=0.7049 by one-way ANOVA) (**Figure 2G**). Given that neurotrophic factors (NFs) are known to promote cell survival, weC hypothesized that withdrawal of NF supplementation from the differentiation medium might reveal a specific vulnerability of mutant MNs, as reported by another group with *C9ORF72* iPSC-derived MNs.^33^ The overall survival of MN cultures decreased by approximately 40% at 6-weeks post-plating when cells were differentiated in medium without NFs compared with complete medium (Isog Ctrl: p=0.0316 by two-way ANOVA) (**Figure 2E**), highlighting the importance of NF supplementation for long-term culture of iPSC-derived MNs. However, viability did not significantly differ between the mutant and control cell lines in absence of NFs, although the viability of TDP-43^A382T^ MNs was not statistically equivalent to control by 6-weeks post-plating (TDP-43^A382T^: p=0.9951; TDP-43^G348C^: p=0.9989 by two-way ANOVA; see **Figure S4D-S4F** for equivalence testing) (**Figure 2E**).

Next, we tested the hypothesis that TDP-43 MNs might be more vulnerable when challenged with cellular stressors, such as glutamate or the oxidative stress inducer ethacrynic acid (EA).^34^ Both glutamate excitotoxicity^35–37^ and oxidative stress^38^ have been proposed to play a role in the pathogenesis of ALS. We first analyzed the viability of MN cultures treated with glutamate for 24 hours (**Figure 2H**). As expected, glutamate treatment induced MN death, as shown by a significant decrease in viability compared with vehicle-treated cultures (Isog Ctrl: p=0.0063; TDP-43^A382T^: p<0.0001; TDP-43^G348C^: p=0.0001 by two-way ANOVA). When comparing the survival of glutamate-treated TDP-43 MNs and isogenic control, we observed a reduced viability in both TDP-43^A382T^ and TDP-43^G348C^ MNs after treatment indicative of an enhanced susceptibility to excitotoxic insults (TDP-43^A382T^: p=0.0093; TDP-43^G348C^: p=0.0056 by two-way ANOVA). Furthermore, TDP-43^A382T^ MNs were more affected by EA treatment than isogenic controls MNs (p=0.0325 by two-way ANOVA). A similar trend was observed in TDP-43^G348C^ MNs, although this effect did not reach statistical significance (p=0.1447 by two-way ANOVA). Taken together, these results indicate that TDP-43 MNs did not display an overt neurodegenerative phenotype at baseline but appear to be more vulnerable to cellular stress.

### Mutant MNs do not accumulate insoluble or phosphorylated TDP-43

TDP-43 aggregates constitute the pathological hallmark of ALS, with a biochemical signature that consists of detergent-insoluble phosphorylated TDP-43 as well as C-terminal fragments of the protein.^4–6^ To assess TDP-43 levels and solubility, we performed protein fractionation of mutant and control MN cultures into total, soluble, and insoluble fractions (**Figure 3A**). Western blot analysis of total lysates showed similar TDP-43 levels between mutant and isogenic control MNs (TDP-43^A382T^: p=0.9997; TDP-43^G348C^: p=0.8636 by one-way ANOVA) (**Figure 3G-3H**). Accordingly, *TARDBP* transcripts levels did not significantly differ between mutant and control samples at all tested timepoints (**Figure S5C**), indicating that the mutations do not cause impairments in the autoregulatory function of TDP-43, where TDP-43 regulate levels of its own *TARDBP* transcript via a negative feedback loop.^39^ When analyzing the soluble (RIPA) and insoluble (urea) protein fractions, mutant and control lysates showed no difference in levels of soluble and insoluble TDP-43 (Soluble fraction TDP-43^A382T^: p=0.7953; TDP-43^G348C^: p=0.9138; Insoluble fraction TDP-43^A382T^: p=0.3321; TDP-43^G348C^: p=0.3823 by one-way ANOVA) (**Figure 3B-3D**), indicating a similar solubility of TDP-43^A382T^ and TDP-43^G348C^ to the wild-type protein. The C-terminal fragment of 35 kDa (CTF-35) was markedly enriched in insoluble fractions, although no differences in insoluble CTF-35 levels were observed in mutant and control MNs (TDP-43^A382T^: p=0.9729; TDP-43^G348C^: p=0.6479 by one-way ANOVA) (**Figure 3E**). Additionally, the CTF-35/TDP-43 ratio did not significantly differ (TDP-43^A382T^: p=0.9266; TDP-43^G348C^: p=0.6185 by one-way ANOVA) (**Figure 3F**), indicating that the TDP-43 variants do not display enhanced C-terminal cleavage compared with control.

**Figure 3.**
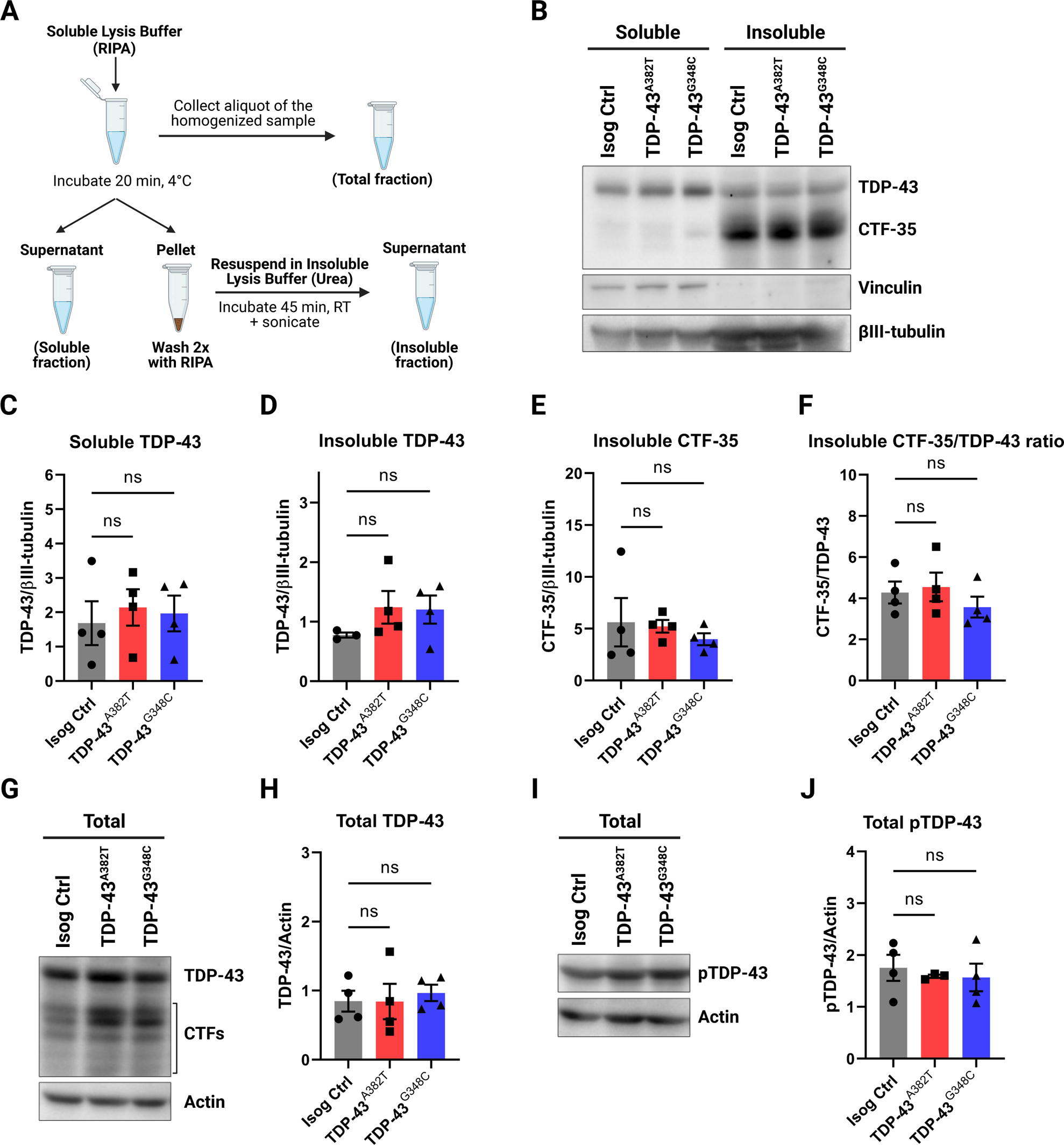
Quantification of TDP-43 levels in total, soluble, and insoluble protein fractions. (A) Schematics representing the fractionation workflow into total (unfractionated), soluble (RIPA), and insoluble (urea) protein fractions. (B-F) Immunoblot (B) and quantification of TDP-43 (C, D) and C-terminal fragment of 35 kDa (CTF-35) (E, F) levels in soluble and insoluble fractions. Vinculin (soluble) and βIII-tubulin were used as fractionation and loading controls, respectively. (G-J) Immunoblot (G, I) and quantification of total levels of TDP-43 (H) and phosphorylated TDP-43 (Ser409/410) (J) in unfractionated lysates. Actin was used as loading control. All data shown as mean ± SEM. Extractions were performed in MNs harvested after 6 weeks post-plating. n=4 independent experiments. See also Figure S5.

Next, we analyzed the phosphorylation state of TDP-43 using an antibody targeting phosphorylated TDP-43 (pTDP-43) (Ser409/410). Using western blotting, we found similar levels of total pTDP-43 in unfractionated lysates of mutant and control MNs (TDP-43^A382T^: p=0.8558; TDP-43^G348C^: p=0.7954 by one-way ANOVA) (**Figure 3I-3J**). Immunostaining showed abundant punctate pTDP-43 in the cytoplasm, with no significant differences in the abundance of pTDP-43^+^ puncta between mutant and control MNs (TDP-43^A382T^: p=0.8600; TDP-43^G348C^: p=0.0926 by one-way ANOVA) (**Figure S5A-S5B**).

### TDP-43^A382T^ and TDP-43^G348C^ do not exhibit changes in nucleocytoplasmic localization

Another prominent feature of TDP-43 pathology is mislocalization of TDP-43 in the cytoplasm. Thus, we performed nuclear/cytosolic protein fractionation experiments to quantify the distribution of TDP-43 (**Figure 4A**). Western blot analysis indicated no significant differences in TDP-43 levels in nuclear and cytosolic fractions of TDP-43 MNs compared with control (Nuclear fraction TDP-43^A382T^: p=0.9882; TDP-43^G348C^: p=0.5749; Cytosolic fraction TDP-43^A382T^: p=0.8879; TDP-43^G348C^: p=0.0534 by one-way ANOVA) (**Figure 4B-4D**). CTF-35 was mainly recovered in cytosolic fractions, at similar levels between TDP-43^A382T^ and control samples (p=0.6863 by one-way ANOVA) (**Figure 4E**). Intriguingly, cytosolic CTF-35 levels appeared to be decreased in TDP-43^G348C^ MNs relative to control (p=0.0472 by one-way ANOVA) (**Figure 4E**). However, the CTF-35/TDP-43 ratio did not significantly differ between samples (TDP-43^A382T^: p=0.9908; TDP-43^G348C^: p=0.9517 by one-way ANOVA) (**Figure 4F**), implying no increased C-terminal cleavage in TDP-43 MNs compared with control in concordance with soluble/insoluble fractionation experiments (**Figure 3F**).

**Figure 4.**
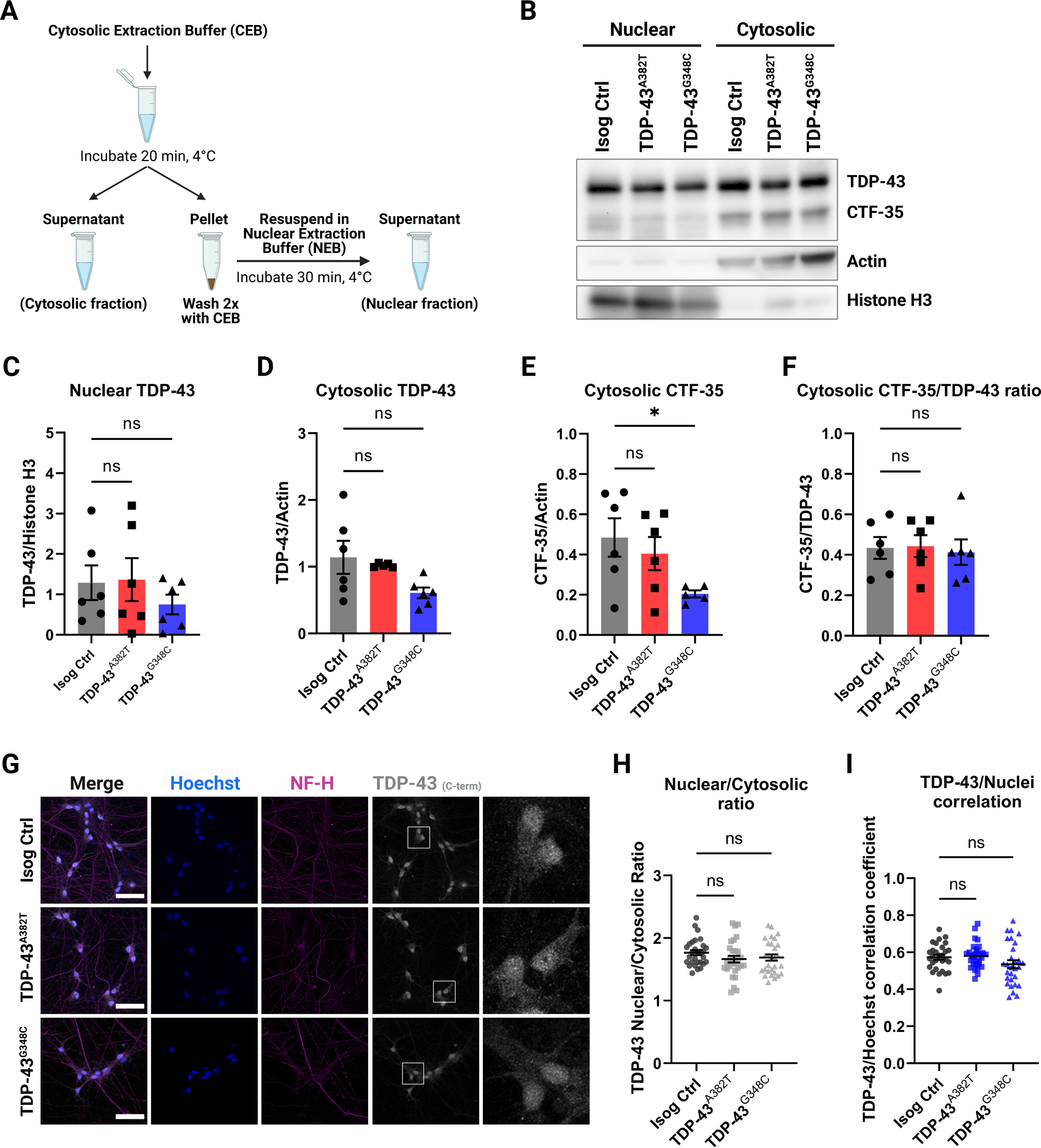
Subcellular distribution of TDP-43 in MNs. (A) Schematics representing the fractionation workflow into nuclear and cytosolic fractions. (B-F) Immunoblot of nuclear and cytosolic fractions (B) and quantification of nuclear (C) and cytosolic (D) TDP-43 levels, and cytosolic C-terminal fragment of 35 kDa (CTF-35) levels (E,F). Histone H3 (nuclear marker) and actin (cytosolic marker) were used as both loading and fractionation controls. n=6 extractions from 4 independent differentiations. Pooled data from MNs harvested 4- and 6-weeks post-plating. (G) Representative images of MNs differentiated for 6 weeks subjected to immunocytochemistry for TDP-43 (C-terminal antibody) and NF-H. Scale bar, 50 µm. (H, I) Quantification of TDP-43 distribution using the nuclear/cytosolic ratio of TDP-43 fluorescence signal intensity (H) and the TDP-43/Hoechst correlation coefficient (I). Individual data points represent per-frame mean values from 5 independent experiments. All data shown as mean ± SEM. See also Figure S6.

To further assess TDP-43 subcellular localization, we performed immunocytochemistry (**Figure 4G**). TDP-43 was predominantly nuclear with some signal detected in the cytoplasm. Immunocytochemical analyses revealed comparable nuclear-to-cytosolic ratios (TDP-43^A382T^: p=0.2261; TDP-43^G348C^: p=0.4055 by one-way ANOVA) (**Figure 4H**) and TDP-43/Hoechst correlation coefficients (TDP-43^A382T^: p=0.9325; TDP-43^G348C^: p=0.1819 by one-way ANOVA) (**Figure 4I**) between TDP-43 MNs and control, indicating similar nucleocytoplasmic localization. These observations were recapitulated with a second TDP-43 antibody targeting the protein’s N-terminus (rather than its C-terminus) (Nuclear-to-cytosolic ratios TDP-43^A382T^: p=0.0501; TDP-43^G348C^: p=0.3535; TDP-43/Hoechst correlation coefficient TDP-43^A382T^: p=0.9637; TDP-43^G348C^: p=0.9996 by one-way ANOVA) (**Figure S6A-S6C**). Compared with C-terminal immunostaining, N-terminal immunostaining showed a more prominent cytosolic signal and TDP-43^+^ puncta could be observed (**Figure S6A**), as previously described.^40^ However, it is worthy noting that TDP-43^+^ puncta were not detected more frequently in mutant MNs compared with control and did not co-localize with pTDP-43^+^ puncta, as would be expected from pathological aggregates. As such, we hypothesize that TDP-43^+^ puncta may represent other condensates such as stress granules, whose assembly have been proposed to be distinct from pTDP-43^+^ aggregate formation.^41–43^ Taken together, these results indicate that mutant MNs do not exhibit TDP-43 pathology as observed in post-mortem tissues.

### Progressive decline in spontaneous neuronal activity in TDP-43 MNs

To assess the activity and functionality of MNs, we performed electrophysiological profiling of MN cultures using multielectrode array (MEA) over a span of 8 weeks post-plating. We observed progressive alteration of spontaneous neuronal activity with prolonged time in culture, as shown by a significant decline in mean firing rate in TDP-43^A382T^ and TDP-43^G348C^ MNs compared with isogenic control MNs by 7-weeks post-plating (TDP-43^A382T^ p=0.0020; TDP-43^G348C^ p=0.0065 by two-way ANOVA) (**Figure 5B**; **Video S2**). Additionally, we noted fewer active electrodes in TDP-43 MN cultures compared with control cultures despite a similar distribution of cells over the electrodes (**Figure 5A** and **5C**), indicating that they are more electrophysiologically silent. Among active electrodes, however, the burst frequency, number of spikes per burst, and burst duration did not significantly differ between TDP-43 and control MN cultures (**Figure S7A-S7C**). Treatment with the sodium-channel blocker tetrodotoxin (TTX) abolished neuronal activity (Isog Ctrl p<0.0001 by two-way ANOVA) (**Figure 5D**), thereby confirming that the recorded signals are due to action potentials and not artifacts.

**Figure 5.**
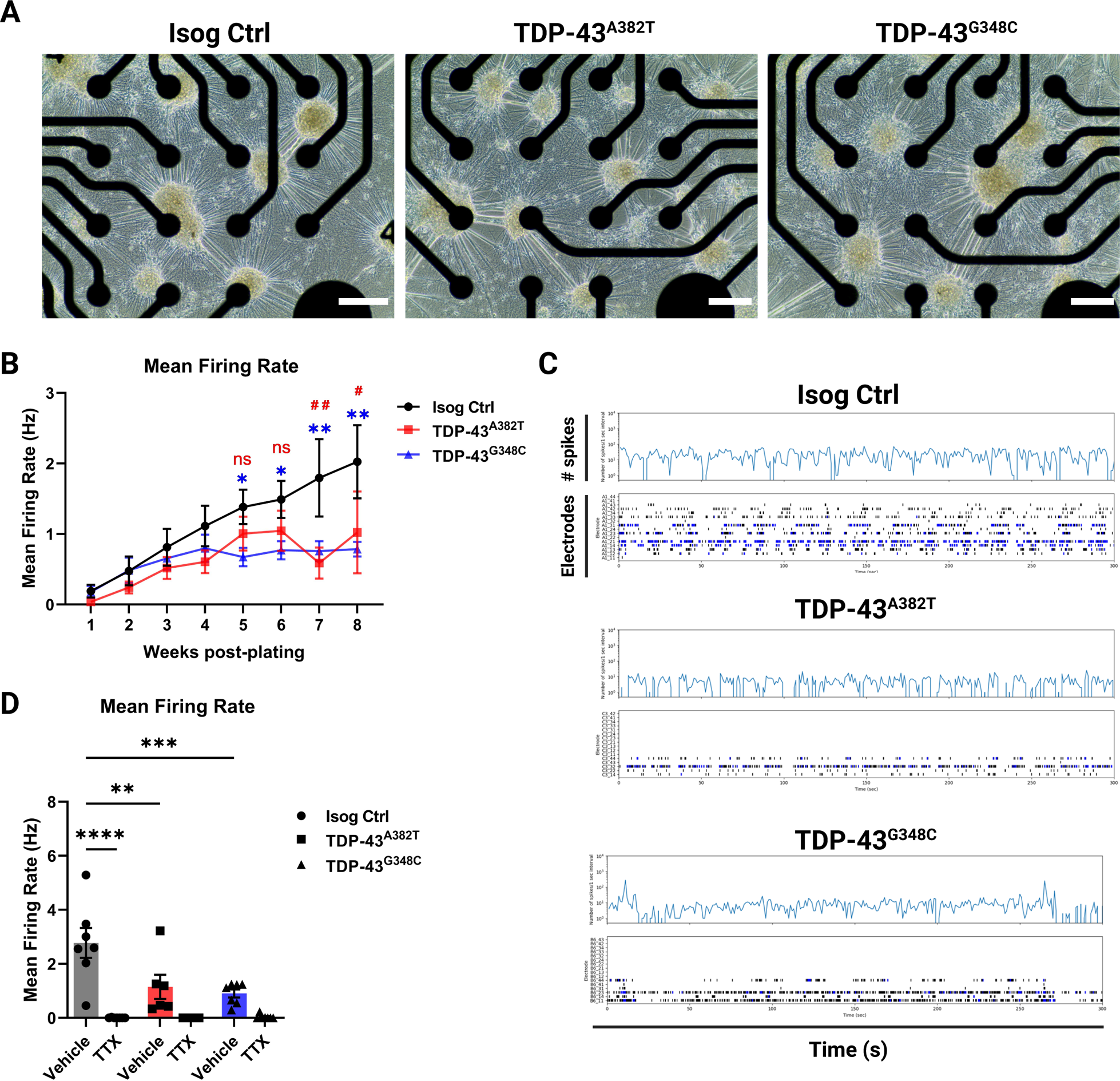
TDP-43 MNs show progressive alterations in spontaneous neuronal activity. (A) Representative phase-contrast images of MNs differentiated for 6 weeks on 24-well MEA plates. Scale bar, 250 μm. (B) Longitudinal changes in mean firing rate of MNs recorded weekly over a span of 8 weeks post-plating. n=11 independent experiments. (C) Spontaneous neuronal activity of MN cultures differentiated for 6 weeks recorded for 300s shown as raster plot and spike histogram. Individual spikes are shown in black and bursts are shown in blue. (D) Effect of TTX treatment on mean firing rate in MNs differentiated for 6 weeks. n=7 independent experiments. All data shown as mean ± SEM. \**p*<0.05, \*\**p*<0.01, \*\*\**p*<0.001, \*\*\*\**p*<0.0001. See also Figure S7.

### TDP-43 MNs exhibit abnormal pre- and post-synaptic puncta

To further study the mechanisms underlying altered neuronal activity, we examined whether TDP-43 MNs would display changes in synapse number and morphology. We performed co-immunostaining for pre-synaptic (synapsin I) and post-synaptic (PSD95) compartments in MN cultures differentiated for 6 weeks (**Figure 6A**) and analyzed mean puncta count, size, and signal intensity. We found no significant change in the number of synapsin I^+^ puncta between TDP-43 MNs and control (TDP-43^A382T^ p=0.2499; TDP-43^G348C^ p=0.8566 by one-way ANOVA) (**Figure 6B**). However, the average size of synapsin I^+^ puncta was moderately increased in TDP-43^A382T^ MNs, but not TDP-43^G348C^ MNs, compared with control (TDP-43^A382T^ p=0.0338; TDP-43^G348C^ p=0.9816 by one-way ANOVA) (**Figure 6C**). We also observed a significant decrease in synapsin I^+^ puncta mean intensity in TDP-43^G348C^ MNs compared with TDP-43^A382T^ and control MNs (TDP-43^A382T^ p=0.4929; TDP-43^G348C^ p=0.0104 by one-way ANOVA) (**Figure 6D**). When analyzing PSD95 immunostaining, the number of PSD95^+^ puncta was significantly decreased in TDP-43^A382T^ MNs and we observed a trend towards fewer PSD95^+^ puncta in TDP-43^G348C^ compared with control (TDP-43^A382T^ p=0.0110; TDP-43^G348C^ p=0.0942 by one-way ANOVA) (**Figure 6E**). Both TDP-43^A382T^ MNs and TDP-43^G348C^ MNs displayed significantly larger PSD95^+^ puncta sizes than control MNs (TDP-43^A382T^ p=0.0009; TDP-43^G348C^ p=0.0007 by one-way ANOVA) (**Figure 6F**). The mean intensity of PSD95^+^ puncta was not significantly different between TDP-43 MNs and control, although a trend towards decreased PSD95^+^ puncta intensity was observed in TDP-43^G348C^ MNs (TDP-43^A382T^ p=0.4348; TDP-43^G348C^ p=0.0522 by one-way ANOVA) (**Figure 6G**). We also analyzed the colocalization of both synapsin I and PSD95 markers. We found no significant change in the number of synapsin I^+^/PSD95^+^ puncta between TDP-43 MNs and control (TDP-43^A382T^ p=0.3545; TDP-43^G348C^ p=0.6667 by one-way ANOVA) (**Figure 6H**). However, the size of synapsin I^+^/PSD95^+^ puncta was significantly larger in TDP-43^A382T^ MNs, but not TDP-43^G348C^ MNs, compared with control (TDP-43^A382T^ p=0.0029; TDP-43^G348C^ p=0.8099 by one-way ANOVA) (**Figure 6I**). Based on these observations, we conclude that TDP-43 MNs exhibit synaptic defects, although these changes appear to vary across the two mutations.

**Figure 6.**
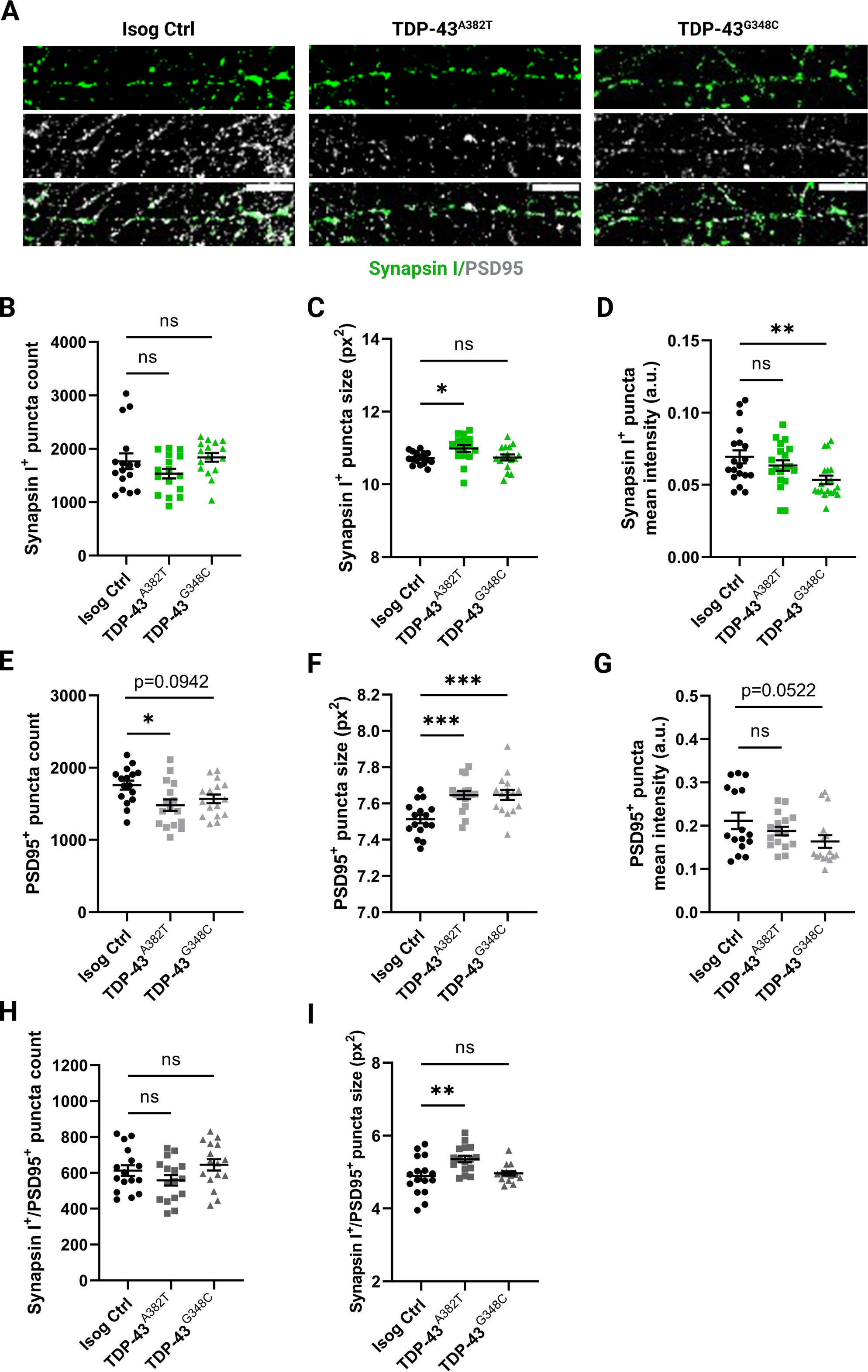
TDP-43 MNs exhibit pre- and postsynaptic abnormalities. (A) Representative images of 6 weeks post-plating MN neurites subjected to immunocytochemistry for synapsin I and PSD95. Scale bar, 10 μm. (B-D) Quantification of the average number (B), size (C), and intensity (D) of synapsin I^+^ puncta. (E-G) Quantification of the average number (E), size (F), and intensity (G) of PSD95^+^ puncta. (H,I) Quantification of the average number (H), and size (I) of synapsin I^+^/PSD95^+^ puncta. Individual points represent per-frame values from 3 independent experiments. All data shown as mean ± SEM. \**p*<0.05, \*\**p*<0.01, \*\*\**p*<0.001.

### TDP-43 variants perturb the expression of synaptic proteins post-transcriptionally

Earlier MEA experiments implied that alterations in activity manifest after prolonged time in culture, leading us to analyze protein levels of presynaptic (synapsin I, synaptophysin) and postsynaptic (PSD95) markers at several timepoints (**Figure 7A**). Western blot analysis indicated increased levels of PSD95 at 2 weeks post-plating in both TDP-43 MN cultures compared with control (TDP-43^A382T^ p=0.0078; TDP-43^G348C^ p=0.0140 by two-way ANOVA), but those levels were not significantly different at other timepoints (**Figure 7B**). We observed similar levels of synaptophysin between samples at all timepoints (**Figure 7D**). Levels of synapsin I, however, were significantly depleted at 6-weeks post-plating, coinciding with the observed decline in mean firing rate (TDP-43^A382T^ p<0.0001; TDP-43^G348C^ p<0.0001 by two-way ANOVA) (**Figure 7C**). We next sought to determine whether dysregulation of synaptic marker expression occurs at the transcriptional level (**Figure 7E-7G**). Despite the prominent decrease in synapsin I protein at 6-weeks post-plating, *SYN1* transcript levels, however, did not significantly differ between TDP-43 and control MNs at this timepoint (TDP-43^A382T^ p=0.9792; TDP-43^G348C^ p=0.9461 by two-way ANOVA) (**Figure 7F**). These results imply that TDP-43 variants perturb the expression of synapsin I post-transcriptionally. In summary, we find that impairments in spontaneous neuronal activity are accompanied by abnormal synaptic marker expression.

**Figure 7.**
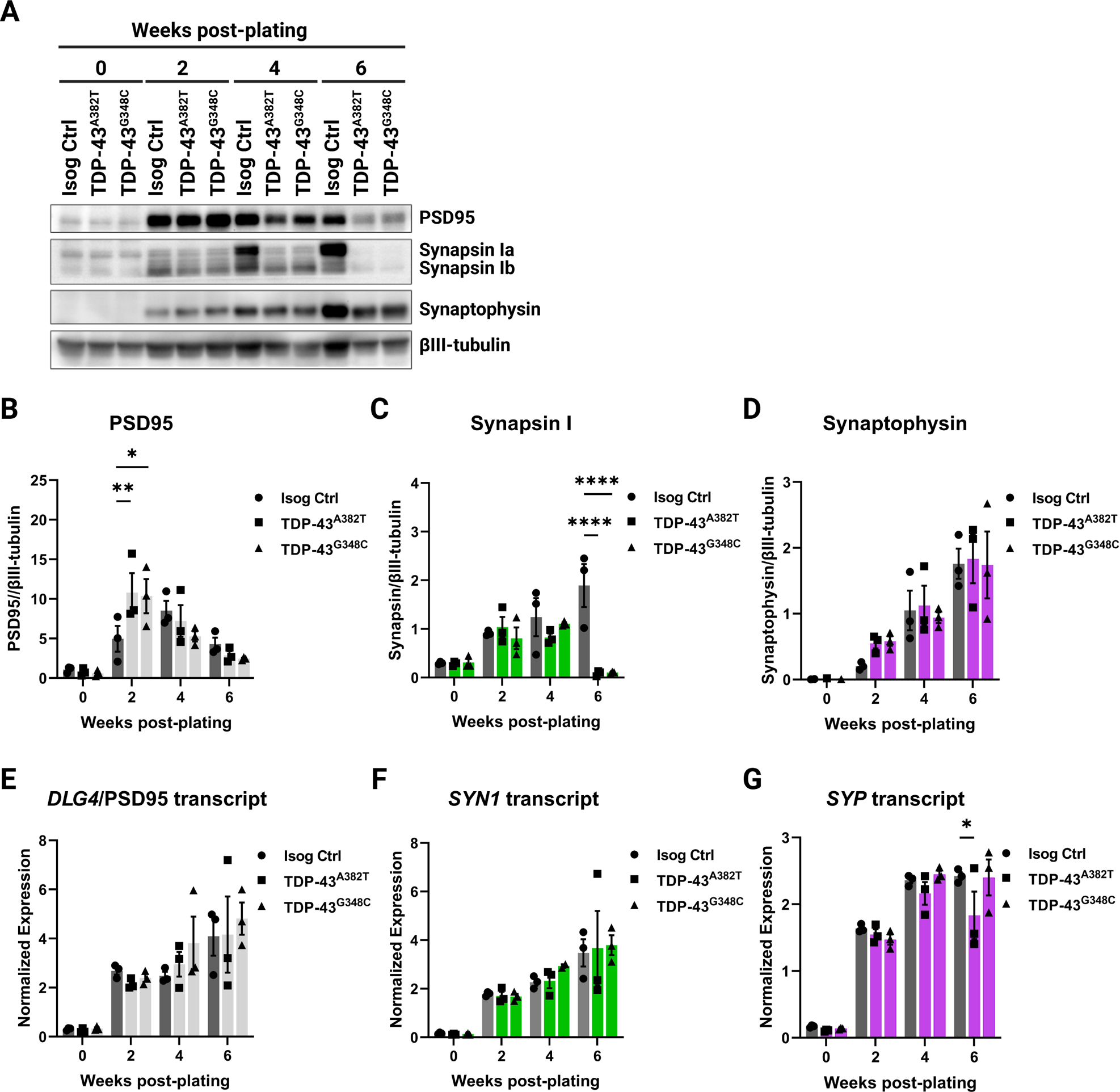
TDP-43 variants lead to decreased synapsin I protein levels but not *SYN1*transcript levels. (A-D) Immunoblot (A) and quantification of protein levels of PSD95 (B), synapsin I (C), and synaptophysin (D) in MNPCs and MNs harvested after 2, 4, and 6 weeks of differentiation. βIII-tubulin was used as loading control. (E-G) Longitudinal quantification of relative transcript levels of *DLG4* (encoding PSD95) (E), *SYN1* (F), and *SYP* (G) using qPCR. All data shown as mean ± SEM. \**p*<0.05, \*\**p*<0.01, \*\*\**p*<0.001, \*\*\*\**p*<0.0001. n=3 independent experiments.

## Discussion

The detection of TDP-43 pathology in almost all ALS cases, along with the identification of disease-causing mutations in the *TARDBP* gene, underscores a central role of TDP-43 dysregulation in ALS pathobiology. Yet, the mechanisms by which *TARDBP* mutations contribute to MN dysfunction and neurodegeneration remains poorly understood. In this study, we harnessed iPSC and CRIPSR/Cas9 gene editing technologies to study the impact of *TARDBP* mutations in a more physiological context. Due to our lack of TDP-43 patient samples, we edited a healthy control iPSC line to generate two homozygous knock-in iPSC lines with *TARDBP* mutations encoding the TDP-43^A382T^ and TDP-43^G348C^ variants, respectively, thereby allowing the comparison of cellular phenotypes between gene-edited cells and their parental cell line. This isogenic experimental design (where cell lines share the same genetic information except for the mutation of interest) is critical to eliminate variability due to genetic background, reprogramming, and differentiation efficiencies, which can affect the reproducibility of experiments in iPSC studies.^44,45^ Furthermore, a knock-in strategy enables the study of these mutations in the homozygous state, which may produce stronger phenotypes than in the heterozygous state. Indeed, the rare ALS patients with homozygous mutations encoding TDP-43^A382T^ have been reported to present with a more severe disease and a complex neurological syndrome in comparison to heterozygous carriers of the same family.^46,47^ We should however point out that, to our knowledge, the mutation encoding TDP-43^G348C^ has never been detected in the homozygous state in patients.

With these edited lines, we tested the hypothesis that MNs differentiated from the *TARDBP* knock-in iPSCs would manifest some features of ALS *in vitro* when cultured for a prolonged period. We found that TDP-43 MNs did not exhibit a cell death phenotype up to 6 weeks of maturation, the latest timepoint investigated. These observations are consistent with earlier reports in which differences in viability between MN cultures derived from *TARDBP* mutant and control iPSC lines were not detected under basal culture conditions.^48–52^ Some studies noted enhanced stress-induced neuronal death following treatment with compounds such as sodium arsenite, staurosporine, MG-132 or LY294002 (a selective PI3K inhibitor),^48,51,53,54^ suggesting some inherent vulnerability conferred by the mutations. Similarly, we observed an increased susceptibility of mutant MNs to glutamate excitotoxicity and, to some extent, to oxidative stress. While we limited our analyses to cell viability post-treatment, stress-based approaches may help induce additional disease-relevant phenotypes and provide valuable insights into the interplay between cellular stress and genetic risk factors.^55^ Notably, an incomplete penetrance of *TARDBP* mutations have been documented^56,57^ (estimated at ∼60% by age 70 in p.A382T carriers^58,59^), suggesting additional factors contributing to disease manifestation.

The absence of overt MN loss (at least under basal conditions) may also reflect the immaturity of MNs at the timepoints examined. In future studies, overcoming the challenges associated with long-term culture of MNs in monolayers (i.e., cell clumping and detachment) or use of three-dimensional (3D) culture models (i.e., spheroids, organoids)^60–62^ can potentially enable neurodegeneration to be observed after several months of culture, without the need for exogenous treatments with stressors to elicit a phenotype. Alternatively, transcription factor-mediated transdifferentiation towards specific neurons, avoiding intermediate proliferative pluripotent stem cell stage, may help retaining specific age-related and epigenetic features involved in neurodegenerative disorders.^63^

As aging is a strong risk factor for ALS, enhancing the maturation of MN cultures may also accelerate the manifestation of end-stage disease features, such as TDP-43 pathology. Here, we show that mutant MNs did not robustly display cytoplasmic mislocalization, aggregation, or accumulation of insoluble TDP-43 under basal conditions. These observations are in line with most recent reports of patient-derived and knock-in iPSC models with *TARBDP* mutations encoding TDP-43^A382T^ or other ALS variants of TDP-43.^41,52,64–70^ Some studies, in contrast, found that mutant MNs recapitulate partial aspects of TDP-43 pathology *in vitro,*^48,53,54,71–73^ sometimes reporting enhanced cytoplasmic distribution of TDP-43 (albeit without nuclear depletion), increased levels of insoluble TDP-43 and lower molecular weight species and/or, in few instances, detection of “preinclusion-like aggregates” by immunocytochemistry or electron microscopy. These discrepancies, together with the absence of overt neurodegeneration, suggest that iPSC-derived MNs may model early stages of ALS.

One prominent finding of the present study is the progressive decline in spontaneous neuronal activity in TDP-43^A382T^ and TDP-43^G348C^ MNs after several weeks in culture. Although we noted abnormal pre- and post-synaptic puncta, we found no significant changes in the number of synapsin I^+^/PSD95^+^ puncta between TDP-43 MNs and control, indicating that altered neuronal activity is not due to a failure of synaptogenesis nor synaptic loss. These results suggest that functional alterations in synaptic activity may arise before the physical disruption of synapses in ALS. Indeed, previous studies of animal models and ALS patients have implicated excitability defects in this disease.^74,75^ Additionally, neuronal hypoexcitability, sometimes preceded by transient early hyperexcitability, has been described in iPSC-derived neurons with mutations in *TARDBP.*^49,51,65^ A possible progression from initial hyper- to hypoexcitability is also supported by studies in ALS mouse models^76–78^ and iPSC-derived models carrying mutations in *C9ORF72,*^49,65,79–81^ *SOD1,*^82–84^ and *FUS,*^82,85^ depending on the timepoint examined. Here, early hyperexcitability was not detected, perhaps due to differences in the electrophysiological method employed.

Accumulating evidence suggests that TDP-43 is involved in synaptic functions, both at central and neuromuscular synapses (reviewed here^86–88^). Pathologically altered TDP-43 has been shown to perturb the expression of synaptic genes in ALS mouse models and patients.^89–92^ Here, we found that synapsin I protein levels, but not *SYN1* transcript levels, were depleted in TDP-43^A382T^ and TDP-43^G348C^ MNs after 6 weeks post-plating, which was coincident with the decline in neuronal activity. These results imply that differences in synapsin I levels result from a post-transcriptional mechanism, such as impairments in mRNA transport, translation and/or mRNA sequestration by TDP-43, as described by several groups.^10,93–95^ Further research will be required to elucidate the molecular mechanisms by which *TARDBP* mutations lead to decreased synapsin I expression, potentially contributing to functional defects.^96,97^ As our analyses were mainly descriptive, this study can not establish a causal link between the hypoactivity phenotype and the synaptic abnormalities observed, which may be explored in future work. Additionally, future studies examining the expression levels of other factors modulating neuronal excitability (e.g., ion channels, glutamate receptor subunits) may provide further insight into the mechanisms underlying the observed impairments in synaptic activity.

Overall, our findings indicate that TDP-43 pathology is not required to induce MN dysfunction and point to early synaptic impairments prior to MN loss in TDP-43-ALS. As the synapse emerges as a promising therapeutic target for ALS, neuronal activity and synapse integrity may serve as disease-relevant phenotypic readouts for drug discovery.

## Limitations of the study

In the present study, we employed a knock-in strategy to study the effects of *TARDBP* mutations in the homozygous state and in isogenic conditions. However, one limitation of this approach is that since the mutant iPSC lines were not derived from patient cells, they may lack ALS genetic modifiers naturally present in patients’ genotypes which could be important contributors to the disease phenotypes. Additionally, experiments were conducted in monolayer cultures enriched in MNs and thus did not take into consideration the potential contribution of other cell types (e.g., astrocytes, microglia, interneurons), which are important areas of interest in ALS research. In particular, astrocytes are known to promote neuronal maturation and synaptogenesis,^98^ and thus astrocyte-MNs co-cultures may be preferable for electrophysiological studies.^80^ Furthermore, monolayer cultures lack cell-to-cell and cell-to-matrix interactions, important cues for cell differentiation and maturation, which may be better recapitulated by 3D models.^99^ Given these limitations, findings with “pure” MN cultures will need to be corroborated in co-culture and 3D models.

## Acknowledgements

We acknowledge Dr. Vincent Soubannier for training and guidance on microscopy image acquisition and analysis, Ghislaine Deyab for guidance on MEA data analysis, Shuming Li for programming the macro for visualization of MEA data, and Dr. Mark R. Aurousseau for his contribution to the development of the glutamate assay. We are also grateful to Dr. Gary A.B. Armstrong for creative discussions and his advice and to Dr. Lenore K. Beitel for proofreading the manuscript. S.L. was supported by the Faculty of Medicine and Health Sciences of McGill University. This work was supported by the Canada First Research Excellence Fund, awarded through the Healthy Brains, Healthy Lives initiative at McGill University; the CQDM FACS program; and the US Department of Defense ALS research program. All figures and schematics were created with BioRender.com.

## Author contributions

Conceptualization, S.L., M.C., and T.M.D.; Methodology, S.L., E.D., M.C., G.M., G.H., and T.M.D.; Software, S.L.; Validation, S.L.; Formal analysis, S.L.; Investigation, S.L., A.N.J., E.D., C.X.-Q.C., N.A., A.K.F.-F., G.H., M.J.C.-M.; Resources, M.C. and T.M.D.; Data curation, S.L.; Writing—original draft, S.L., M.C., and T.M.D.; Writing—review and editing, S.L., E.D., M.J.C.-M., G.M., M.C., and T.M.D.; Visualization, S.L., M.C. and T.M.D.; Supervision, M.C. and T.M.D.; Project Administration, S.L., M.C., and T.M.D.; Funding acquisition, T.M.D. All authors have read and agreed to the published version of the manuscript.

## STAR Methods

### RESOURCE AVAILABILITIES

#### Lead contact

Further information and requests should be directed to the lead contact, Thomas M. Durcan.

#### Materials availability

Cell lines generated in this study will be made available on request, under the open science framework of the Neuro, and through a cost recovery model.

#### Data and code availability

Derived data supporting the findings of this study are available from the lead contact upon reasonable request.

### EXPERIMENTAL MODEL AND SUBJECT DETAILS

#### iPSC lines and culture

The use of human cells in this study was approved by McGill University Health Center Research Ethics Board (DURCAN_iPSC / 2019-5374). To knock-in selected mutations, we used the previously characterized control cell line AIW002-02, reprogrammed from peripheral blood mononuclear cells (PBMCs) of a 37-year-old Caucasian male, as previously described.^26^ A summary of the iPSC lines used can be found in **Table S1**. iPSCs were maintained on dishes coated with Matrigel (Corning Millipore; Cat#354277) in mTeSR1 (StemCell Technologies; Cat#85850) and passaged at 80% confluence using Gentle Cell Dissociation Reagent (StemCell Technologies; Cat#07174). Cultures were routinely tested for mycoplasma using the MycoAlert Mycoplasma Detection kit (Lonza; Cat#LT07-318).

#### Differentiation of iPSCs into MNs

Differentiation of iPSCs into MNs was performed using a previously published protocol.^25^ Briefly, iPSCs were plated onto Matrigel-coated T25 flasks in “neural induction medium”, composed of basic neural medium (1:1 mixture of DMEM/F12 medium (Gibco; Cat#10565–018) and Neurobasal medium (Life Technologies; Cat#21103–049), 0.5X N2 (Life Technologies; Cat#17502–048), 0.5X B27 (Life Technologies; Cat#17504–044), 0.5X GlutaMAX (Gibco, Cat#35050-061), 1X antibiotic-antimycotic (Gibco, Cat#15240–062), and 100 µM ascorbic acid (Sigma-Aldrich; Cat#A5960)) supplemented with 2 µM SB431542 (Selleckchem; Cat#S1067), 2 µM DMH1 (Selleckchem; Cat#S7146), 3 µM CHIR99021 (Selleckchem; Cat#S2924), with medium fully changed every other day. At Day 6, cells were split onto 10 µg/ml Poly-L-ornithine (PLO) (Sigma-Aldrich; Cat#P3655) and 5 µg/ml laminin (Sigma-Aldrich; Cat#L2020)-coated T75 flasks at a 1:3 to 1:6 ratio in “patterning medium” (basic neural medium supplemented with 1 µM CHIR99021, 2 µM SB431542, 2 µM DMH1, 0.1 µM retinoic acid (RA, Sigma-Aldrich; Cat#R2625) and 0.5 µM purmorphamine (PMN, Sigma-Aldrich; Cat#SML-0868)), with medium fully changed every other day. At Day 12, cells were passaged at a ratio of 1:3 to 1:6 onto PLO/laminin-coated T75 flasks and cultured in “expansion medium” (basic neural medium supplemented with 3 µM CHIR99021, 2 µM SB431542, 2 µM DMH1, 0.1 µM RA and 0.5 µM PMN, and 0.5 mM valproic acid (VPA, Sigma-Aldrich; Cat#P4543)), with medium fully changed every other day for 6 days. MNPCs were then cryopreserved for later use or passaged and maintained in the expansion medium.

To perform final differentiation into MNs, MNPCs were dissociated as single cells using Accutase and plated onto PLO/laminin (Life Technologies; Cat#23017-015)-coated dishes in “final differentiation medium” (basic neural medium supplemented with 0.5 µM RA, 0.1 µM PMN, 0.1 µM Compound E (CpdE, StemCell Technologies; Cat#73954), and 10 ng/ml insulin-like growth factor-1 (IGF-1, Peprotech; Cat#100–11), brain-derived neurotrophic factor (BDNF, Peprotech; Cat#450–02) and ciliary neurotrophic factor (CNTF, Peprotech; Cat#450–13)), with weekly half-changes. Alternatively, to decrease cell clumping, MNPCs were first passaged in “priming medium” (basic neural medium supplemented with 0.5 µM RA and 0.1 µM PMN)) for 6 days with medium changed every other day, before plating in final differentiation medium. Whenever cells were dissociated for passaging or plating throughout the differentiation protocol from iPSCs to MNs, the culture medium was supplemented with 10 µM ROCK inhibitor Y-27632 (Selleckchem; Cat#S1049) for the first 24 h to improve survival.

### METHOD DETAILS

#### CRISPR/Cas9 genome editing and validation

To genetically edit *TARDBP*, CRISPR reagents were transfected into iPSCs using the P3 Primary Cell 4D Nucleofector™ X Kit S (Lonza; Cat#V4XP-3032), as previously described.^25^ Briefly, iPSCs at 50% confluency were dissociated with Accutase (StemCell Technologies; Cat#07922) and 500,000 cells were resuspended in 25 μL of Cas9: sgRNA ribonucleoprotein (RNP)-ssODN-buffer mix, consisting of 1 µL of Cas9 protein (stock 61 µM), 3 µL of sgRNA (stock 100 µM), and 1 µL of ssODNs (stock 100 µM) in 20 µL of nucleofection buffer P3. The reaction mixture was then electroporated using the CA137 program in a Nucleofector 4D device (Lonza; Cat#AAF-1002B). The sequences of the sgRNAs and ssODNs used are provided in **Table S2**.

After limiting dilution, gene-edited clones were identified by ddPCR (QX200™ Droplet Reader; Bio-Rad; Cat#1864003). The detection of the modified nucleotide by ddPCR was based on a TaqMan® assay including two PCR primers and two DNA probes fused with different fluorophores (FAM and HEX), with one probe specific to the original allele and the other probe to the edited allele. Locked Nucleic Acid (LNA®) probes were designed following the manufacturer’s criteria. Sequence integrity of successful clones was assessed using Sanger sequencing. The sequences of the primers and probes used for ddPCR, and Sanger sequencing are provided in **Table S3**. Knock-in iPSCs underwent quality control testing, which included karyotyping, genomic stability analysis, and STR profiling.

#### Karyotyping and genomic stability analysis

DNA was extracted from iPSCs using a Genomic DNA Mini Kit (Geneaid; Cat#GB100). Genomic stability was assessed using the hPSC Genetic Analysis Kit (StemCell Technologies; Cat#07550) according to the manufacturer’s instructions. Reactions were run on a QuantStudio 5 Real-Time PCR system (Applied Biosystems; Cat#A28140). The copy number of a control region in chr 4p was used for normalization. G-band karyotyping was performed on 50-60% confluent iPSCs cultured for 72 h at The Centre for Applied Genomics of The Hospital for Sick Children (Toronto, ON).

#### STR profiling

DNA was extracted from iPSCs using a Genomic DNA Mini Kit (Geneaid; Cat#GB100). STR profiling was performed at the Centre for Applied Genomics of The Hospital for Sick Children (Toronto, ON) using the the GenePrint® 10 System. This system allows co-amplification and detection of nine human loci, including the ASN-0002 loci (TH01, TPOX, vWA, Amelogenin, CSF1PO, D16S539, D7S820, D13S317 and D5S818) as well as D21S11.

#### Quantitative PCR

Total RNA was isolated with the miRNeasy kit (Qiagen; Cat#217004) with DNase treatment (Qiagen; Cat#79256) following the manufacturer’s instructions. cDNA synthesis was performed using 500 ng of RNA with the M-MLV Reverse Transcriptase kit (ThermoFischer; Cat#28025013) in a total volume of 40 μL. Real-time qPCR reactions were set up in triplicates with the TaqMan® Fast Advanced Master Mix (Applied Biosystems; Cat#A44360) and TaqMan® Assays (ThermoFischer; Cat#4331182) and run on a QuantStudio 5 Real-Time PCR system (Applied Biosystems; Cat#A28140). The geometric mean of *ACTB* and *GAPDH* was used for normalization. The following TaqMan® probes were used: *ACTB* (Hs01060665_g1), *GAPDH* (Hs02786624_g1), *NES* (Hs04187831_g1), *PAX6* (Hs01088114_m1), *OLIG2* (Hs00377820_m1), *MNX1* (HB9) (Hs00907365_m1), *ISL1* (Hs00158126_m1), *CHAT* (Hs00758143_m1), *SLC18A3* (VAChT) (Hs00268179_s1), *LHX3* (Hs01033412_m1), *FOXP1* (Hs00212860_m1), *TARDBP* (Hs00606522_m1), *DLG4* (PSD95) (Hs01555373_m1), *SYN1* (Hs00199577_m1), *SYP* (Hs00300531_m1) and are listed in **Table S4**.

#### Immunofluorescence staining

Cells were fixed in 4% formaldehyde for 20 min at room temperature and washed three times with PBS. After fixation, cells were permeabilized with 0.2% Triton X-100 in PBS for 10 min and blocked in 5% normal donkey serum (NDS, Millipore; Cat#S30-100), 1% bovine serum albumin (BSA, Multicell; Cat#800–095-CG) and 0.05% Triton X-100 in PBS for 1 h at room temperature. Cells were then incubated in primary antibodies diluted in blocking solution overnight at 4°C, washed three times with PBS, and incubated with secondary antibodies for 2 h in the dark at room temperature. Cells were then washed in PBS and counterstained with Hoechst 33342 DNA dye (Life Technologies; Cat#H3570) diluted 1:2000 in PBS for 5 min. Coverslips were mounted with Aqua-Poly/Mount (Polysciences; Cat#18606–5). The following primary antibodies were used: rabbit anti-Nanog (Abcam; Cat#ab21624; 1:200), mouse anti-TRA-1-60 (Stem Cell Tech; Cat#60064; 1:200), mouse anti-SSEA-4 (Santa Cruz; Cat# sc-21704; 1:200), goat anti-Oct-3/4 (Santa Cruz; Cat#sc-8628; 1:500), mouse anti-PAX6 (DSHB; Cat#PAX6; 1:100), rabbit anti-OLIG2 (Millipore; Cat#AB9610; 1:100), rabbit anti-Nestin (Abcam; Cat#ab92391; 1:250), mouse anti-HB9 (DSHB; Cat#81.5C10-c; 1:50), mouse anti-ISL1/2 (DSHB; Cat#39.4D5-c; 1:50), goat anti-ChAT (Millipore; Cat#MAP144P; 1:50), rabbit anti-VAChT (Sigma-Aldrich; Cat#SAB4200559; 1:100), chicken anti-NF-H (EnCor Biotech; Cat#CPCA-NF-H; 1:1000), mouse anti-βIII-tubulin (Millipore; Cat#MAB5564; 1:1000), rabbit anti-TDP-43 (C-terminal) (Proteintech; Cat#12892-1-AP; 1:2000), rabbit anti-TDP-43 (N-terminal) (Proteintech; Cat#10782-2-AP; 1:1000), mouse anti-pTDP-43 (Ser409/410) (Proteintech; Cat#66318-1-lg; 1:500), rabbit anti-Synapsin (Calbiochem; Cat#57477; 1:500), mouse anti-PSD95 (Millipore; Cat#MABN68; 1:250). The following secondary antibodies were used: Donkey anti-Mouse IgG DyLight® 488 (Abcam; Cat#ab96875), Donkey anti-Mouse IgG DyLight® 550 (Abcam; Cat#ab96876), Donkey anti-Mouse IgG DyLight® 650 (Abcam; Cat#ab96878), Donkey anti-Rabbit IgG DyLight® 488 (Abcam; Cat#ab96891), Donkey anti-Rabbit IgG DyLight® 650 (Abcam; Cat#ab96894), Donkey anti-Goat IgG Alexa Fluor® 488 (Jackson Immunoresearch; Cat#705-545-147), Donkey anti-Goat IgG Alexa Fluor® 647 (Invitrogen; Cat#A21447), Donkey anti-Chicken Alexa Fluor® 647 (Jackson Immunoresearch; Cat#703-605-155). All antibodies used are listed in the **Key resources table**.

#### Microscopy and image acquisition

Phase-contrast images were acquired using an EVOS XL Core (Thermo Fischer Scientific). The time-lapse movie depicting differentiation of MNPCs into MNs was generated using the JuLI^TM^ Stage system and software (NanoEntek). For immunostainings of iPSC and MNPC markers, wide-field fluorescence microscopy images were acquired using an automated EVOS FL-Auto2 imaging system (Thermo Fischer Scientific) and a ZEISS Axio Observer Z1 microscope, respectively, with consistent exposure time across conditions. Confocal microscopy images were acquired for TDP-43, synaptic, and MN markers immunostainings using a Leica TCS SP8 microscope, with consistent gain and laser settings across conditions.

#### Viability assay

MNPCs were plated as 15,000 cells per well in opaque white optical 96-well plates coated with PLO/laminin. Cells were cultured in final differentiation medium for one week, after which they were treated with 1 μM cytosine arabinoside (AraC, Sigma-Aldrich; Cat#C6645) overnight (∼17 h) to eliminate any residual proliferating cells. The next day, medium was fully changed to final differentiation medium with or without supplementation neurotrophic factors (i.e., BDNF, CNTF, and IGF-1), with weekly half-changes until initiation of the assay. Breathe-Easy sealing membranes (Sigma-Aldrich) were applied onto the plates to minimize evaporation. The ATP-based luminescence assay Cell-Titer Glo® (Promega; Cat#G7570) was used to determine the viability of cultures at multiple timepoints (1-, 2-, 4-, and 6-weeks post-plating), following the manufacturer’s instructions. The luminescence readings were acquired using a GloMax Microplate Reader (Promega). The percentage of viability was determined by normalizing the raw luminescence values to the 1-week reading (where viability was assumed to be 100%) of each cell line to account for differences in plating.

For the glutamate and oxidative stress assays, MNPCs were plated as described above and cultured in complete differentiation medium for 4 weeks, after which the assay was initiated. For the glutamate assay, half of the medium was removed and replaced by medium supplemented with glutamate (Sigma-Aldrich; Cat#G1626) and cyclothiazide (CTZ, Tocris; Cat#0713) for a final concentration of 0.1 mM glutamate and 50 μM CTZ for 24 h. The vehicle treatment condition consisted of an equivalent concentration of CTZ without glutamate. For oxidative stress assays, MNs were treated with ethacrynic acid (EA, Sigma-Aldrich; Cat#SML1083) at a final concentration of 50 μM or with vehicle (DMSO) and were incubated overnight (17 h). For all conditions, the percentage of viability was determined by normalizing the raw luminescence values to those of untreated wells of each cell line.

#### Soluble/insoluble protein fractionation

MNs differentiated for 4 and 6 weeks were fractionated into total, soluble, and insoluble protein fractions. MN cultures were harvested using Accutase and centrifuged for 3 min at 1,300 g. Cells were washed by resuspending the cell pellet with PBS, transferred into 1.5 mL microcentrifuge tubes, and centrifuged for 5 min at 5,000 g. Cell pellets were resuspended in 10 packed cell volume (pcv) of ice-cold “soluble” lysis buffer (RIPA buffer (Millipore; Cat#20-188) supplemented with protease inhibitors (Roche; Cat#11697498001) and phosphatase inhibitors (Roche; Cat#04906837001). For example, for a pcv of 40 μL, 400 μL of RIPA buffer were used. The cell homogenate was vortexed before and after a 30-min incubation at 4°C on a rotator, then an aliquot of the homogenate (the total fraction) was collected in a new microtube. The rest of the homogenate was centrifuged at 10,000 g for 10 min. The supernatant (the soluble fraction) was collected in a new microtube, while the pellet (containing RIPA-insoluble proteins and cell debris) was washed twice by resuspending the pellet in 250 μL RIPA buffer, rotating for 5 min at 4°C and centrifuging at 5,000 g for 5 min at 4°C. The washes (supernatants) were discarded, and the pellet resuspended in “insoluble” lysis buffer (7 M urea, 2 M thiourea, 4% CHAPS, 0.03 M Tris-HCl pH 8.5, supplemented with protease and phosphatase inhibitors). The volume of urea buffer used was one quarter of the RIPA buffer volume. Then, the lysate was incubated for 45 min on a shaker at room temperature and sonicated with a probe sonicator (3 pulses of 10 s, 40% amplitude). After centrifugation at 10,000 g for 20 min, the supernatant (the insoluble fraction) was collected in a new microtube. All protein extracts were stored at −80°C until use.

#### Nuclear/cytosolic protein fractionation

MNs differentiated for 6 weeks were fractionated into nuclear and cytosolic fractions based on a previously published protocol,^72^ with a few modifications. MN cultures were harvested using Accutase and centrifuged for 3 min at 1,300 g. Cells were washed by resuspending the cell pellet with PBS, transferred into 1.5 mL microtubes, and centrifuged for 5 min at 5,000 g. Cell pellets were resuspended by pipetting up and down gently with 10 pcv of ice-cold Cytosolic Extraction Buffer (CEB) (50 mM Tris–HCl pH 6.5, 100 mM NaCl, 300 mM Sucrose, 3 mM MgCl2, 0.15% NP40, 4 mM DTT, 40 mM EDTA, supplemented with protease and phosphatase inhibitors). Cell homogenates were incubated at 4°C for 20 min on a shaker, then centrifugated for 5 min at 5,000 g at 4°C. The supernatant (the cytosolic fraction) was transferred into a new microtube and centrifuged again for 10 min at 10,000 g at 4°C to eliminate residual cell debris, while the pellet (containing nuclei) was washed twice by resuspension in 250 μL CEB, incubation for 5 min at 4°C on a shaker and centrifugation at 5,000 g for 5 min at 4°C. The washes (supernatants) were discarded, and the pellet was resuspended in ice-cold Nuclear Extraction Buffer (NEB) (RIPA buffer supplemented with protease and phosphatase inhibitors). The volume of RIPA lysis buffer used was one quarter of the CEB volume used. Then, the lysate was incubated for 30 min at 4°C on a shaker, centrifuged at 10,000 g for 10 min at 4°C, and the supernatant (the nuclear fraction) was collected in a new microtube. All protein extracts were stored at −80°C until use.

#### Western blotting

MNs were fractionated into total (unfractionated), soluble, insoluble, nuclear, and cytosolic fractions as described above. Protein concentrations of soluble, nuclear, and cytosolic fractions were determined by using the DC Protein Assay (Bio-Rad; Cat#5000111). A total of 20 μg of protein per sample in a final volume of 20 μL in Laemmli buffer was resolved by 7.5% or 10% SDS/PAGE and transferred to PVDF or nitrocellulose membranes using a Trans-Blot Turbo Transfer System (Bio-Rad), except for nuclear and cytosolic fractions where 8 μg of protein per sample was used. For total (unfractionated) and insoluble fraction (urea) samples, an equivalent volume of the correspondent RIPA-soluble counterparts was used for sample preparation.^100^ After transfer, membranes were blocked in 5% BSA or 5% milk in TBS-T 0.1% for 1 h at room temperature and incubated with primary antibodies in blocking solution overnight at 4°C. After three washes with TBS-T 0.1%, membranes were incubated with horseradish peroxidase (HRP)-conjugated secondary antibodies (1:10000) in blocking solution for 2 hours at room temperature. Blots were developed with Clarity Western ECL Substrate or Clarity Max Western ECL Substrate (Bio-Rad; Cat#170-5061; Cat#1705062) and pictures were acquired with a ChemiDoc MP Imaging System (Bio-Rad). Semiquantitative analysis of western blots was performed with the Image Lab 6.0.1 software (Bio-Rad), using as loading controls βIII-tubulin (for soluble and insoluble fractions), Histone H3 (for nuclear fractions) and actin (for total and cytosolic fractions). The following primary antibodies were used: chicken anti-NF-H (Abcam; Cat#ab4680; 1:5000), mouse anti-βIII-tubulin (Millipore; Cat#MAB5564; 1:20000), rabbit anti-Cleaved Caspase 3 (Cell Signaling; Cat#9661; 1:500), rabbit anti-TDP-43 (C-terminal) (Proteintech; Cat#12892-1-AP; 1:1000-1:4000), mouse anti-pTDP-43 (Ser409/410) (Proteintech; Cat#66318-1-lg; 1:1000), rabbit anti-Vinculin (Abcam; Cat#ab129002; 1:1000), rabbit anti-Histone H3 (Cell Signaling; Cat#4499; Clone D1H2; 1:2000), mouse anti-Actin (Milipore; Cat#MAB1501; Clone C4; 1:20000), rabbit anti-Synapsin (Calbiochem; Cat#57477; 1:500), mouse anti-PSD95 (Millipore; Cat#MABN68; 1:500), mouse anti-Synaptophysin (Sigma-Aldrich, Cat#S5768, 1:500). The following secondary antibodies were used: HPR-conjugated anti-Mouse (Jackson Immunoresearch, Cat#115-035-003), HPR-conjugated anti-Rabbit (Jackson Immunoresearch, Cat#111-035-144), HPR-conjugated anti-Chicken (Jackson Immunoresearch, Cat#703-035-155). All antibodies used are listed in the **Key resources table**.

#### MEA recording

A total of 50,000-100,000 MNPCs per well were plated as droplets in differentiation medium onto the electrode area of Cytoview MEA 24-well plates (Axion Biosystems) coated with PLO/laminin. Cells were allowed to adhere to the electrode area in the incubator for 1 hour, after which 0.5 mL of differentiation medium was added to each well. On the day of the recording, artificial cerebrospinal fluid (aCSF) was freshly prepared from a 10X stock solution as previously described.^60^ Sterile-filtered aCSF was added to MN cultures, and the plate was returned to the incubator for at least 1 hour before the recording. Next, the plate was transferred into a Maestro Edge MEA system (Axion Biosystems) and was allowed to equilibrate in the machine for 5 min. Spontaneous neuronal activity was recorded for 5 min using the AxIS Navigator 1.5.1.12 software (Axion Biosystems). Recordings were performed weekly starting at 1-week post-plating. To ensure that the recorded MEA signals are not artifacts, MN cultures differentiated for 6 weeks were treated with vehicle (H_2_O) or TTX (Sigma-Aldrich; Cat#T8024) at a final concentration of 1 µM, after which recordings were immediately performed.

Phase-contrast images of each well were acquired weekly using a EVOS XL Core (Thermo Fisher Scientific). Individual electrode recordings were manually excluded if MNs were detached and/or if proliferative cellular contaminants were overlying the electrode. If ≥50% of the electrodes of a well met these exclusion criteria, the entire well recording was excluded from the analysis. The macro for visualization of MEA data into raster plots and spike histograms is available at https://github.com/dxe303/MiCM-summer-project.

### QUANTIFICATION AND STATISTICAL ANALYSIS

#### Image analysis

All image analyses were performed with CellProfiler 4.0.7 (freely available from https://cellprofiler.org/).^101^ For analysis of MNPC and MN markers showing a predominantly nuclear staining (i.e., PAX6, OLIG2, HB9, ISL1/2), images were segmented to identify (i) Hoechst-stained nuclei and (ii) cells showing immunoreactivity for the marker using the “IdentifyPrimaryObjects” module. The total counts of both object types were used to calculate a percentage of cells positive for each marker. For analysis of MNPC and MN markers showing a predominantly cytoplasmic/axonal staining (i.e., Nestin, ChAT, VAChT), the total area was determined using the “Threshold” module and was normalized to the total nuclei count.

For NF-H and βIII-tubulin immunostainings, the axonal network analysis was based on a previously published method.^102^ Briefly, “IdentifyPrimaryObjects” was used to identify Hoechst-stained nuclei and cell bodies were identified using “DilateObjects”. After image pre-processing steps with “Smooth” and “EnhanceOrSuppressFeatures” modules, axonal networks were detected using “Threshold”. After subtracting cell bodies using “MaskImage”, total axonal area was determined using “MeasureImageAreaOccupied”. Axonal branching was determined using “MorphologicalSkeleton” and “MeasureObjectSkeleton” modules.

For analysis of TDP-43 subcellular localization, Hoechst images were used to identify nuclei using “IdentifyPrimaryObjects” and TDP-43 images were used to identify cell bodies using “IdentifySecondaryObjects”. Nuclei were subtracted from cell bodies to determine the cytoplasmic regions using “IdentifyTertiaryObjects”. The mean TDP-43 intensity in nuclear and cytoplasmic regions were determined using “MeasureObjectIntensity” and were used to calculate the nuclear-to-cytosolic ratio for TDP-43 immunofluorescence. The per-image correlation coefficients between Hoechst and TDP-43 images were determined using the “MeasureColocalization” module.

For quantification of pTDP-43^+^ puncta, Hoechst images were used to identify nuclei using “IdentifyPrimaryObjects” and pTDP-43 images were used to identify cell bodies using “IdentifySecondaryObjects”. Image pre-processing was performed using “GaussianFilter” and “EnhanceOrSuppressFeatures” modules (Feature type: Speckle). pTDP-43^+^ puncta were identified within cell bodies using “MaskObjects” to subtract axons and background signal, followed by “IdentifyPrimaryObjects”. The per-frame puncta count was normalized to the number of nuclei.

For quantification of synaptic puncta, z-stack confocal images of neurons co-stained for synapsin I (pre-synaptic), PSD95 (post-synaptic), and ChAT (motor neuron) were acquired. Single plane 2D images underwent pre-processing using the “GaussianFilter” module followed by the “EnhanceOrSuppressFeatures” module (Feature type: Speckle). Pre- and postsynaptic puncta were identified with “IdentifyPrimaryObjects” using Otsu’s thresholding method. Advanced settings were optimized to filter out dim puncta (background) and to distinguish clumped objects. Double positive puncta were identified using “MaskObjects” to keep the overlapping regions between synapsin I and PSD95 objects. Puncta mean intensity and size were determined using “MeasureObjectIntensity” and “MeasureObjectSizeShape”, respectively.

#### Statistical tests

Biological replicates were defined as independent differentiations unless otherwise specified. Grubbs’ test was used to determine significant outliers. Statistical analyses were performed with the GraphPad Prism 9.3.0 software. Data distribution was assumed to be normal although this was not formally tested. Differences between multiple groups were analyzed using one-way or two-way analysis of variance (ANOVA) tests. Means and standard errors of the mean were used for data presentation. Significance was defined as *p*<0.05.

Equivalence testing was performed on MN marker expression and viability data collected from the control and mutant lines using 90% confidence intervals (CIs) as described elsewhere.^103,104^ For each comparison, the 90% CI of the mean difference and the effect size of Cohen’s *d* were calculated using the following web app: https://www.psychometrica.de/effect_size.html (option 2). The lower (*Δ_L_*) and upper (*Δ_U_*) equivalence bounds (which define equivalent and non-equivalent groups) were empirically defined for our data set, adjusted to our sample sizes and degrees of freedom of the two-group comparison tests. When the two-group mean difference (A-B) is positive and for an effect size Cohen’s *d* close to −0.8, *ΔL* is expected to be close to the 90% CI lower value. When the two-group mean difference (A-B) is negative, and for an effect size Cohen’s *d* close to 0.8, *ΔU* is expected to be close to the 90% CI upper value. Using 90% CIs and Cohen’s *d* calculated from our data set, we estimated these values to be −1.8 and 1.8, respectively. We plotted the 90% CIs of mean differences for each comparison (**Figure S4**) and then rejected the hypothesis of equivalence between two groups when (i) the lower value of the 90% CI of mean difference was below −1.8 (*ΔL*) or when (ii) the upper value of the 90% CI mean difference was above 1.8 (*ΔU*).

## Video legends

**Video S1. Differentiation of MNPCs into MNs. Related to Figure 1**. Time-lapse movie depicting the differentiation of MNPCs into MNs during 2 weeks post-plating. Scale bar, 150 μm.

**Video S2. Electrophysiological recording of MN cultures using MEA. Related to Figure 5**. Movie depicting spontaneous neuronal activity in MNs differentiated for 7 weeks using the AxIS Navigator 1.5.1.12 software (Axion Biosystems). Warmer colors indicate greater changes in local field potentials.

## Web References

ALSoD database. https://alsod.ac.uk/. Accessed 10 April 2023. Related publications: https://doi.org/10.1002/humu.22157, https://doi.org/10.3109/17482968.2011.584629

