## Supplemental Information for "Homozygous ALS-linked mutations in TARDBP/TDP-43 lead to hypoactivity and synaptic abnormalities in human iPSC-derived motor neurons"

### Table of Contents


#### I. Supplemental Figures

Figure S1. Validation of CRISPR/Cas9 gene editing by ddPCR

Figure S2. Characterization of *TARDBP* knock-in iPSCs

Figure S3. Characterization of iPSC-derived MNPCs and MNs

Figure S4. Equivalence testing with confidence intervals

Figure S5. Mutant MNs do not accumulate detergent-insoluble or phosphorylated TDP-43

Figure S6. TDP-43 variants do not exhibit changes in nucleocytoplasmic localization

Figure S7. Supplemental neuronal activity measurements recorded using multielectrode array

#### II. Supplemental Tables

Table S1. Overview of iPSC lines used

Table S2. Sequences of sgRNAs and ssODNs used in making of *TARDBP* knock-in iPSC lines

Table S3. List of primers and affinity probes used for ddPCR or Sanger sequencing

Table S4. List of TaqMan probes

### I. Supplemental Figures

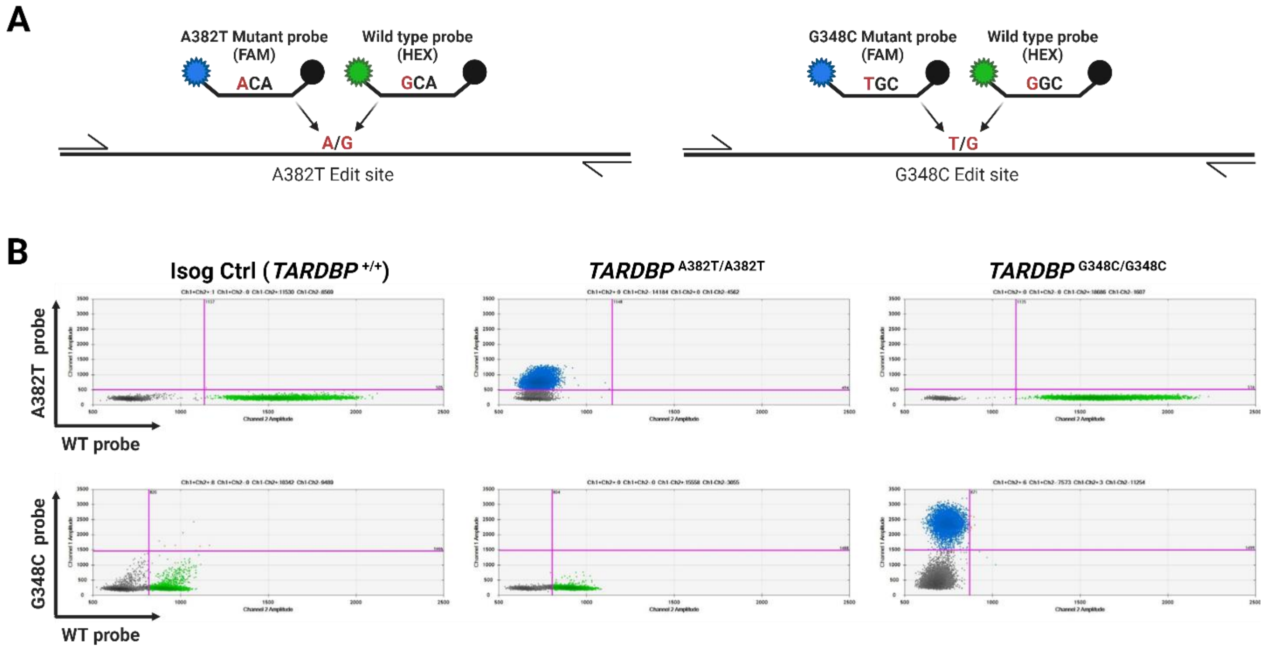


**Figure S1. Validation of CRISPR/Cas9 gene editing by ddPCR. Related to Figure 1.**

(A and B) Pairs of mutants (FAM, blue) and wild-type (HEX, green) probes designed to target the edited or wild-type alleles, respectively (A). ddPCR scatter plots confirming correct gene editing and homozygosity of iPSC lines (B).

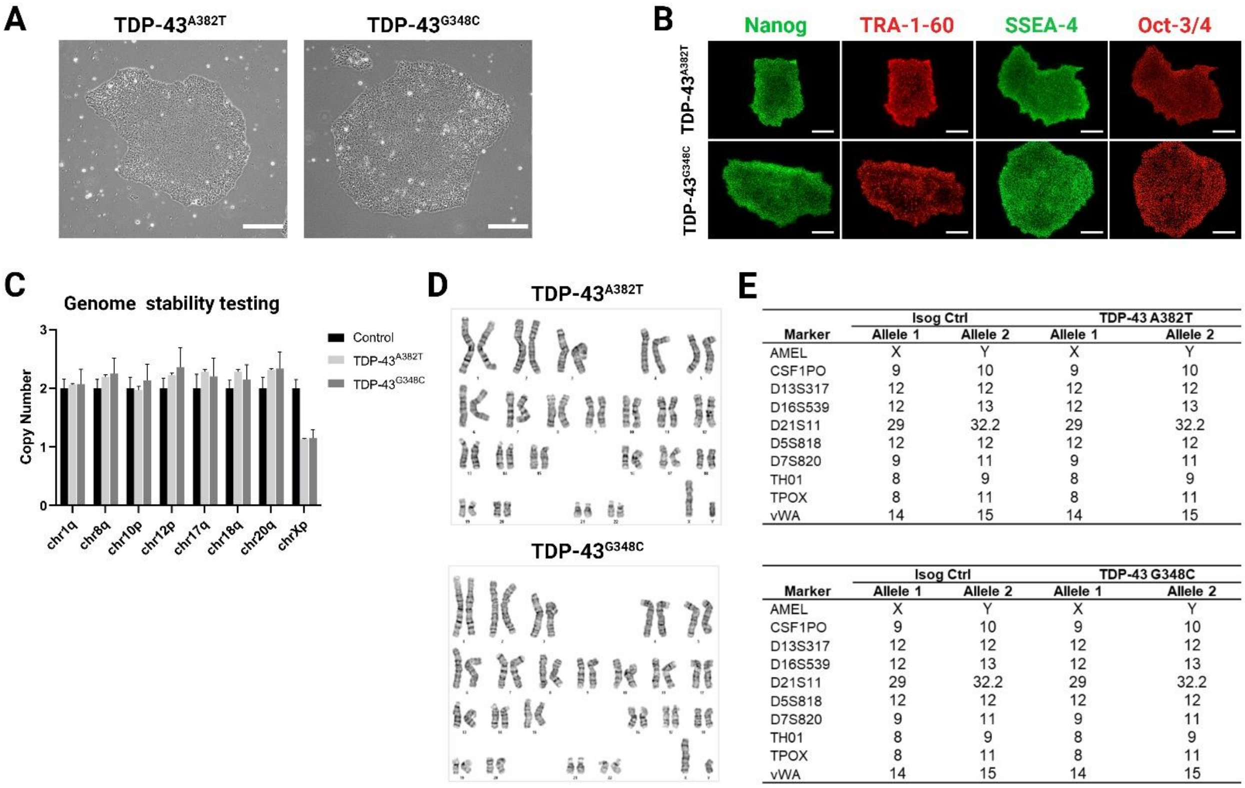


| Marker | Isog Ctrl Allele 1 | Isog Ctrl Allele 2 | TDP-43 A382T Allele 1 | TDP-43 A382T Allele 2 |
|---|---|---|---|---|
| AMEL | X | Y | X | Y |
| CSF1PO | 9 | 10 | 9 | 10 |
| D13S317 | 12 | 12 | 12 | 12 |
| D16S539 | 12 | 13 | 12 | 13 |
| D21S11 | 29 | 32.2 | 29 | 32.2 |
| D5S818 | 12 | 12 | 12 | 12 |
| D7S820 | 9 | 11 | 9 | 11 |
| TH01 | 8 | 9 | 8 | 9 |
| TPOX | 8 | 11 | 8 | 11 |
| vWA | 14 | 15 | 14 | 15 |

| Marker | Isog Ctrl Allele 1 | Isog Ctrl Allele 2 | TDP-43 G348C Allele 1 | TDP-43 G348C Allele 2 |
|---|---|---|---|---|
| AMEL | X | Y | X | Y |
| CSF1PO | 9 | 10 | 9 | 10 |
| D13S317 | 12 | 12 | 12 | 12 |
| D16S539 | 12 | 13 | 12 | 13 |
| D21S11 | 29 | 32.2 | 29 | 32.2 |
| D5S818 | 12 | 12 | 12 | 12 |
| D7S820 | 9 | 11 | 9 | 11 |
| TH01 | 8 | 9 | 8 | 9 |
| TPOX | 8 | 11 | 8 | 11 |
| vWA | 14 | 15 | 14 | 15 |

**Figure S2. Characterization of *TARDBP* knock-in iPSCs. Related to Figure 1.**

(A) Representative phase-contrast images of *TARDBP* knock-in iPSCs. Scale bar, 250 μm.
(B) Representative images of *TARDBP* knock-in iPSCs subjected to immunocytochemistry for pluripotency-associated markers Nanog, TRA1-60, SSEA-4, and OCT-3/4. Scale bar, 250 μm.
(C,D) Genomic stability analyses. *TARDBP* knock-in iPSC lines have normal chromosome copy numbers, as assessed by qPCR. Data shown as mean ± SEM of technical triplicates. The control used here was provided by the manufacturer of the genome stability testing kit (C). Edited iPSC lines display normal G-band karyotypes (D).
(E) STR analysis confirming the isogeneity of *TARDBP* knock-in iPSCs with the parental control line (AIW002-02).

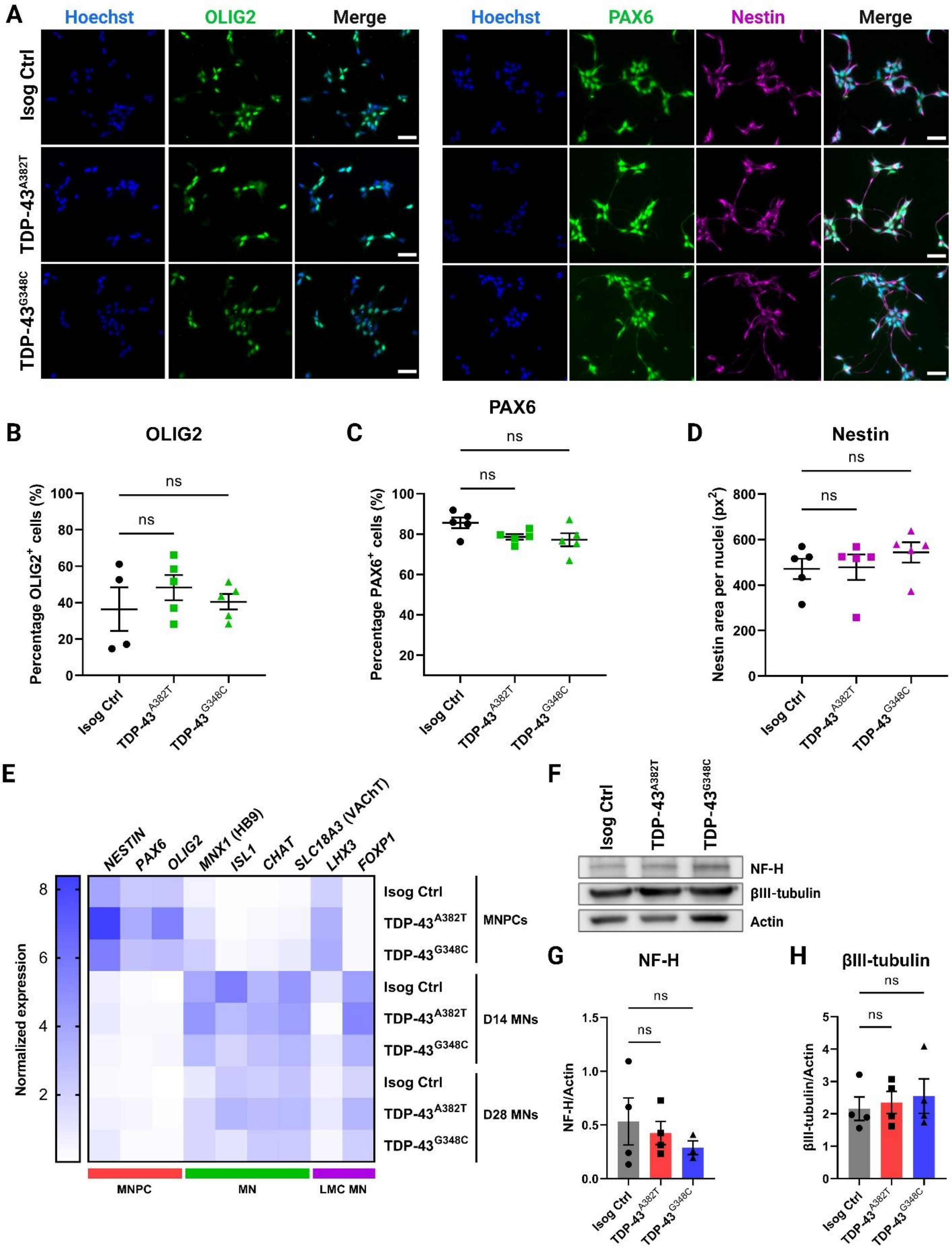

A
Hoechst
OLIG2
Merge
Hoechst
PAX6
Nestin
Merge
Isog Ctrl
TDP-43A382T
TDP-43G348C
B
OLIG2
Percentage OLIG2+ cells (%)
ns
C
PAX6
Percentage PAX6+ cells (%)
D
Nestin
Nestin area per nuclei (px2)
E
NESTIN
PAX6
OLIG2
MNX1 (HB9)
ISL1
CHAT
SLC18A3 (VAChT)
LHX3
FOXP1
Normalized expression
MNPCs
D14 MNs
D28 MNs
MNPC
MN
LMC MN
F
NF-H
βIII-tubulin
Actin
G
NF-H
NF-H/Actin
H
βIII-tubulin
βIII-tubulin/Actin

**Figure S3. Characterization of iPSC-derived MNPCs and MNs. Related to Figures 1 and 2.**

(A-D) Representative images (A) and quantification of MNPCs differentiated from iPSCs subjected to immunocytochemistry for common MNPC markers OLIG2 (B), PAX6 (C), and Nestin (D). Scale bar, 100 μm. n=5 replicates from at least 2 independent inductions from iPSCs. Data shown as mean ± SEM.
(E) qPCR heatmap showing normalized transcripts levels of MNPC (red), MN (green), and LMC (magenta) markers during differentiation of MNPCs into MNs. Mean plotted. n=3 independent experiments.
(F-H) Immunoblot (F) and quantification of total levels of neurofilament heavy (NF-H) (G) and βIII-tubulin (H). Actin was used as loading control. Extractions were performed in MNs harvested after 6 weeks post-plating. n=4 independent experiments. Data shown as mean ± SEM.

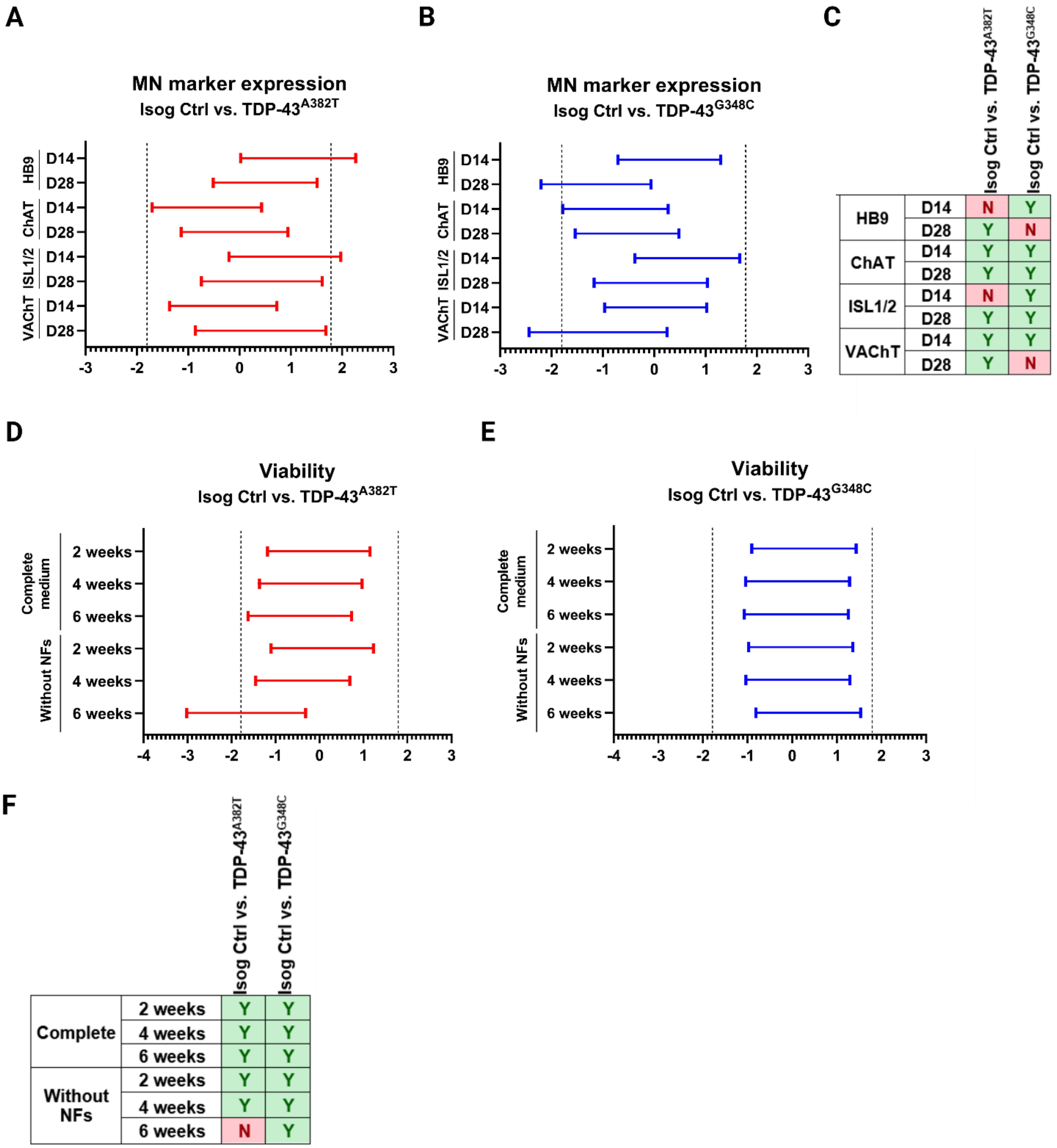


C

| | | Isog Ctrl vs. TDP-43A382T | Isog Ctrl vs. TDP-43G348C |
|---|---|---|---|
| HB9 | D14 | N | Y |
| | D28 | Y | N |
| ChAT | D14 | Y | Y |
| | D28 | Y | Y |
| ISL1/2 | D14 | N | Y |
| | D28 | Y | Y |
| VAChT | D14 | Y | Y |
| | D28 | Y | N |

F

| | | Isog Ctrl vs. TDP-43A382T | Isog Ctrl vs. TDP-43G348C |
|---|---|---|---|
| Complete | 2 weeks | Y | Y |
| | 4 weeks | Y | Y |
| | 6 weeks | Y | Y |
| Without NFs | 2 weeks | Y | Y |
| | 4 weeks | Y | Y |
| | 6 weeks | N | Y |

**Figure S4. Equivalence testing with confidence intervals. Related to Figures 1 and 2.**

(A-C) 90% confidence intervals with equivalence bounds $\Delta_L$ = -1.8 and $\Delta_U$ = 1.8 for comparisons of MN markers expression by immunocytochemistry in TDP-43$^{A382T}$ MNs (A) or TDP-43$^{G348C}$ MNs (B) compared with isogenic control MNs after 2-weeks (D14) and 4-weeks (D28) of final differentiation, with results summary (Equivalence Yes/No) (C).

(E-F) 90% confidence intervals with equivalence bounds $\Delta_L$ = -1.8 and $\Delta_U$ = 1.8 for comparisons of MN viability in TDP-43$^{A382T}$ MNs (E) or TDP-43$^{G348C}$ MNs (F) compared with isogenic control MNs at 2-. 4- and

6-weeks of final differentiation in medium with or without neurotrophic factors (NF) supplementation, with results summary (Equivalence Yes/No) (F).

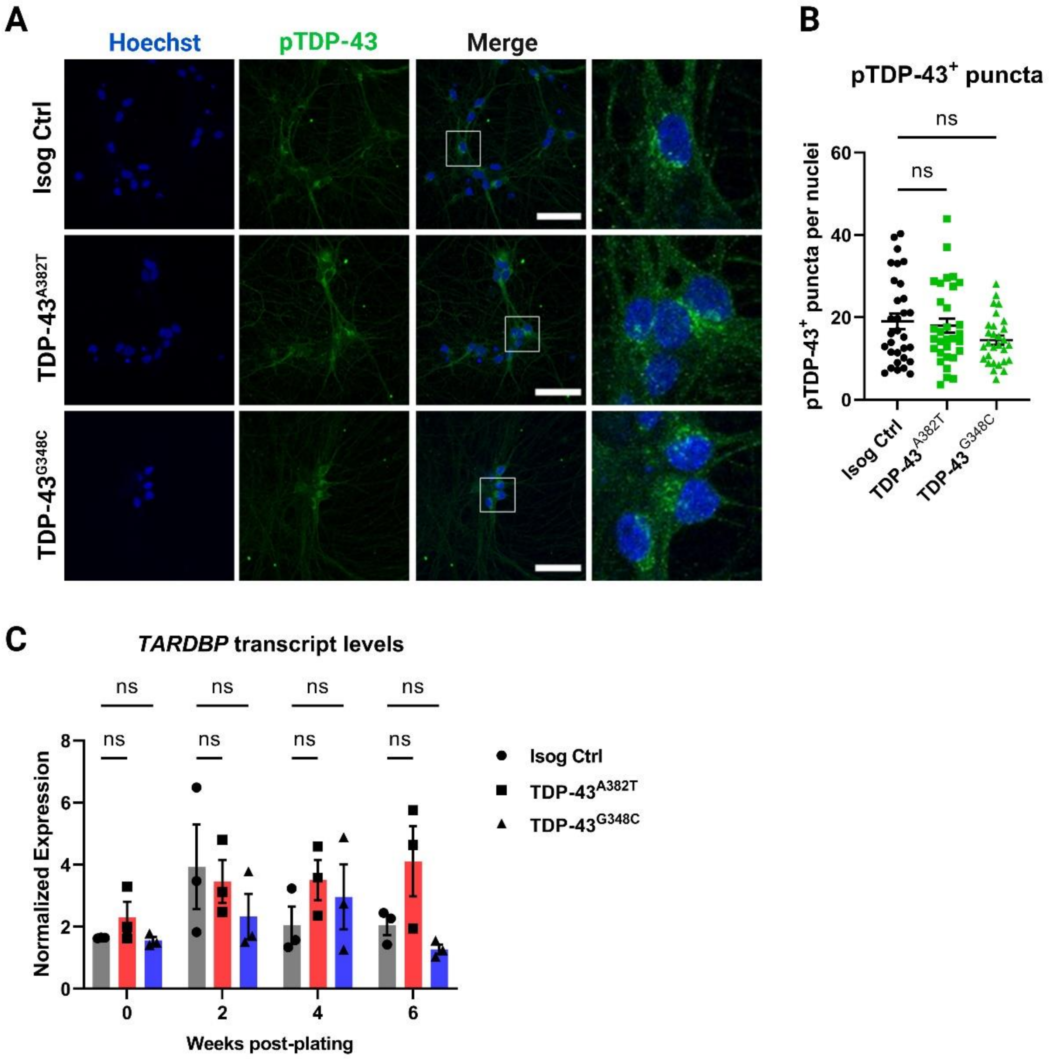


**Figure S5. Mutant MNs do not accumulate *TARDBP* transcripts or phosphorylated TDP-43. Related to Figure 3.**

(A) Representative images MNs differentiated for 6 weeks subjected to immunocytochemistry for phosphorylated TDP-43 (Ser409/410) showing punctate cytosolic staining. Scale bar, 50 µm.
(B) Quantification of pTDP-43$^+$ puncta. Individual data points represent per-frame mean values from 5 independent experiments.
(C) Relative transcript levels of *TARDBP* at several timepoints determine by qPCR. n=3 independent experiments.
All data shown as mean ± SEM.

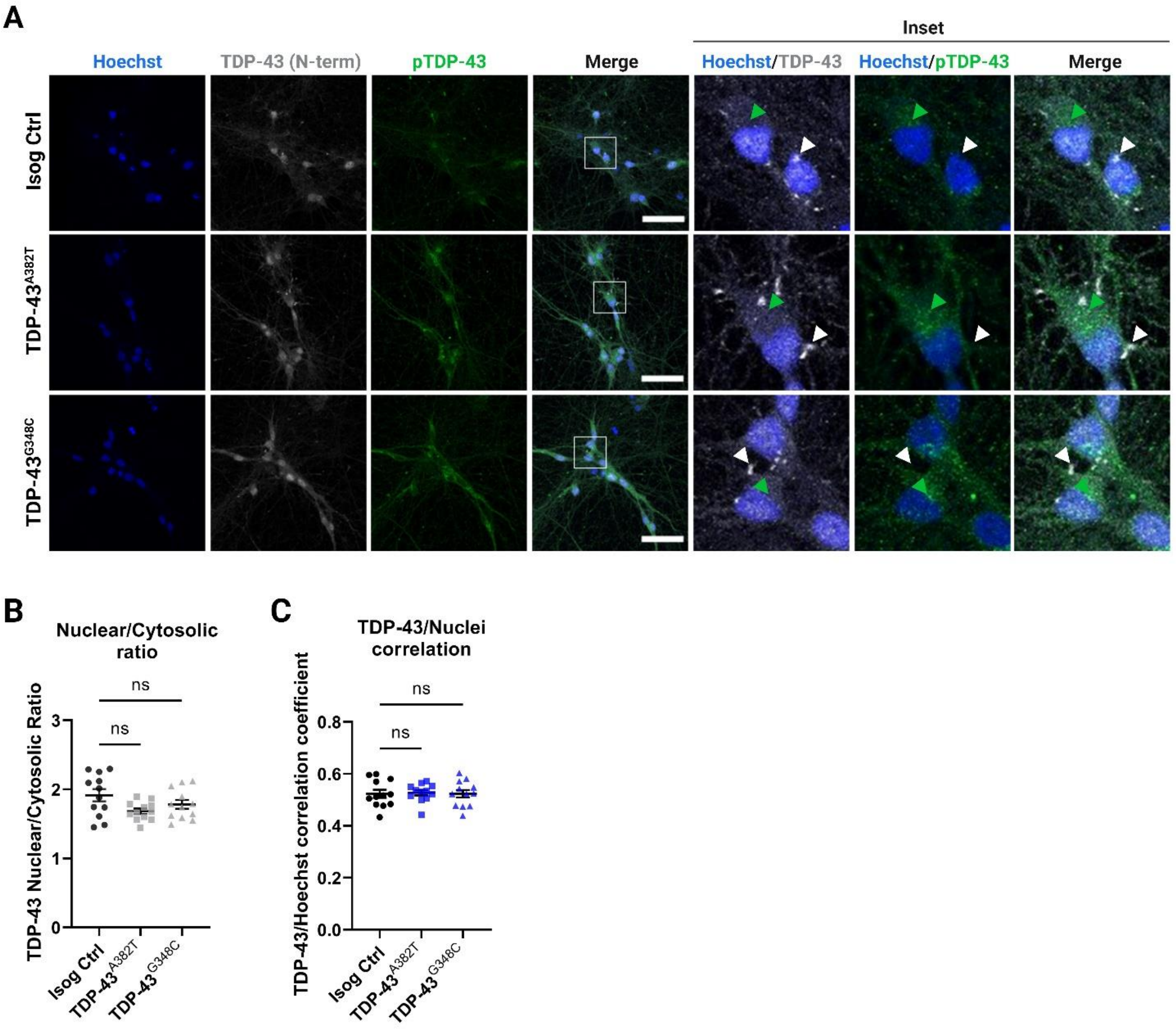


**Figure S6. TDP-43 variants do not exhibit changes in nucleocytoplasmic localization.** Related to **Figure 4.**

(A) Representative immunostainings of MNs differentiated for 6 weeks with an antibody targeted to the N-terminus of TDP-43 showing TDP-43 subcellular distribution and cytosolic TDP-43$^{+}$ puncta. TDP-43$^{+}$ puncta (white arrows) do not colocalize with pTDP-43$^{+}$ puncta (green arrows). Scale bar, 50 µm.

(B,C) Quantification of TDP-43 distribution using the nuclear/cytosolic ratio of TDP-43 fluorescence signal intensity (B) and the TDP-43/Hoechst correlation coefficient (C). Individual data points represent per-frame mean values from 2 independent experiments. Data shown as mean ± SEM.

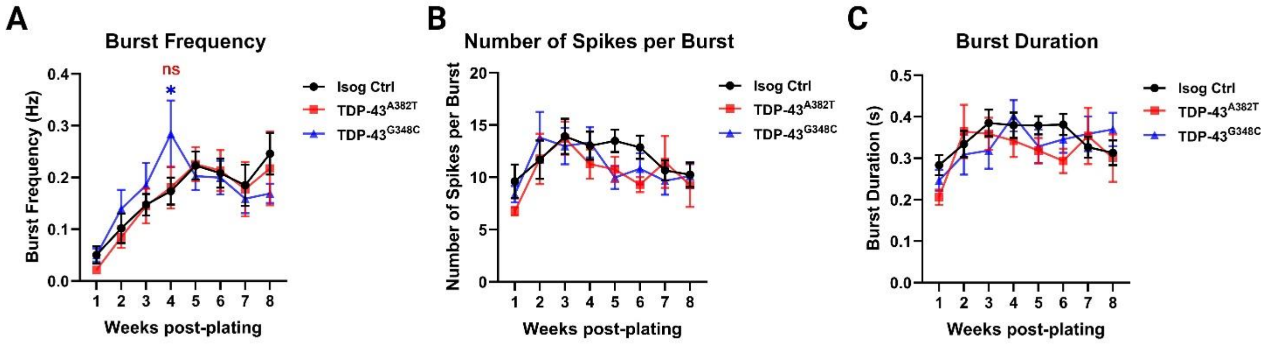


**Figure S7. Supplemental neuronal activity measurements recorded using multielectrode array. Related to Figure 5.**

(A-C) Longitudinal changes in burst frequency (A), number of spikes per burst (B), and burst duration (C) of MN cultures recorded weekly over a span of 8 weeks. n=11 independent experiments. All data shown as mean ± SEM. *$p$<0.05.

#### Supplemental Tables

##### Table S1. Overview of iPSC lines used

| Cell line ID | ALS mutation | Sex | Age | Ethnicity | Primary cell line | Reprogramming method |
|---|---|---|---|---|---|---|
| AIW002-02 | None | Male | 37 | Caucasian | PBMC | Sendai virus |
| *TARDBP* A382T/AIW002-02 | p.A382T | Male | 37 | Caucasian | Knock-in | n/a |
| *TARDBP* G348C/AIW002-02 | p.G348C | Male | 37 | Caucasian | Knock-in | n/a |

* PBMC, peripheral blood mononuclear cell; n/a, not applicable.

##### Table S2. Sequences of sgRNAs and ssODNs used in making of *TARDBP* knock-in iPSC lines

| Cell line ID | gRNA | ssODN template |
|---|---|---|
| *TARDBP* A382T/AIW002-02 | UCUAAUUCUGGUGCAGCAAU | GGCCTTCGGTTCTGGAAATAACTCTTATAGTGGCTCT AATTCTGGTGCAACAATCGGTTGGGGATCAGCATCC AATGCAGGGTCGGGCAGTGGTTTTAATGG |
| *TARDBP* G348C/AIW002-02 | GCCAGCCAGCAGAACCAGUC | GAGCAGTTGGGGTATGATGGGCATGTTAGCCAGCC AGCAGAACCAGTCATGCCCATCGGGTAATAACCAAA ACCAAGGCAACATGCAGAGGGAGCCAAACCAGG |

##### Table S3. List of primers and affinity probes used for ddPCR or Sanger sequencing

| | ***TARDBP* A382T/AIW002-02** | ***TARDBP* G348C/AIW002-02** |
|---|---|---|
| probe-HEX (WT)* | ATT+G+C+TG+CA+CC | AGTC+A+G+GC+CC |
| probe-FAM (Mutant)* | ATT+G+T+TG+C+AC+CA | AG+TC+A+T+GC+CCAT |
| ddPCR primer-F | CTTCGGTTCTGGAAATAACTCTTATAG | TTAGCCAGCCAGCAGAA |
| ddPCR primer-R | CCCAGCCAGAAGACTTAGAA | GTTATTTCCAGAACCGAAGGC |
| Sanger seq primer-F | GCTTTGGGAATCAGGGTGGA | GCTTTGGGAATCAGGGTGGA |
| Sanger seq primer-R | ACTCCACACTGAACAAACCA | ACTCCACACTGAACAAACCA |

* "+" signs in front of nucleotides indicate the location of Locked Nucleic Acids (LNA®).

**Table S4. List of TaqMan probes**

| Gene | Reference |
|---|---|
| ACTB | Hs01060665_g1 |
| GAPDH | Hs02786624_g1 |
| NES | Hs04187831_g1 |
| PAX6 | Hs01088114_m1 |
| OLIG2 | Hs00377820_m1 |
| MNX1/HB9 | Hs00907365_m1 |
| ISL1 | Hs00158126_m1 |
| CHAT | Hs00758143_m1 |
| SLC18A3/VAChT | Hs00268179_s1 |
| LHX3 | Hs01033412_m1 |
| FOXP1 | Hs00212860_m1 |
| TARDBP | Hs00606522_m1 |
| DLG4/PSD95 | Hs01555373_m1 |
| SYN1 | Hs00199577_m1 |
| SYP | Hs00300531_m1 |
